# Dietary αKG inhibits SARS CoV-2 infection and rescues inflamed lungs to restore normal O_2_ saturation in animals

**DOI:** 10.1101/2022.04.02.486853

**Authors:** Sakshi Agarwal, Simrandeep Kaur, Tejeswara Rao Asuru, Garima Joshi, Nishith M Shrimali, Anamika Singh, Oinam Ningthemmani Singh, Puneet Srivastva, Tripti Shrivastava, Sudhanshu Vrati, Milan Surjit, Prasenjit Guchhait

## Abstract

Our recent works described the rescue effect of α-ketoglutarate (αKG, a metabolite of Krebs cycle) on thrombosis and inflammation in animals. αKG augments activity of prolyl hydroxylase 2 (PHD2), which in turn degrades proline residues of substrates like phosphorylated Akt (pAkt) and hypoxia inducible factor (HIF)α. Here we describe the inhibitory effect of octyl αKG on pAkt as well as on HIF1α/HIF2α, and in turn decreasing SARS CoV-2 replication in Vero E6 cells. αKG failed to inhibit the viral replication and Akt phosphorylation in PHD2-knockdown U937 cells transiently expressing ACE2. Contrastingly, triciribine (TCN, an Akt-inhibitor) inhibited viral replication alongside a downmodulation of pAkt in PHD2-KD cells. Dietary αKG significantly inhibited viral infection and rescued hamsters from thrombus formation and inflammation in lungs, the known causes of acute respiratory distress syndrome (ARDS) in COVID-19. αKG supplementation also reduced the apoptotic death of lung tissues in infected animals, alongside a downmodulation of pAkt and HIF2α. αKG supplementation neither affected IgG levels against SARS CoV-2 RBD protein nor altered the neutralization antibody response against SARS CoV-2. It did not interfere with the percentage of interferon-γ positive (IFNγ+) CD4+ and IFNγ+CD8+ T cells in infected animals. The extended work in balb/c mice transiently expressing ACE2 showed a similar effect of αKG in reducing accumulation of inflammatory immune cells and cytokines, including IL6, IL1β and TNFα, in lungs as well as in circulation of infected animals. Pro-thrombotic markers like platelet microparticles and platelet-leukocyte aggregates were reduced significantly in infected mice after αKG supplementation. Importantly, αKG supplementation restored the O_2_ saturation (SpO_2_) in circulation of SARS CoV-2 infected hamsters and mice, suggesting a potential therapeutic role of this metabolite in COVID-19 treatment.

## INTRODUCTION

Akt, also known as protein kinase B, regulates many crucial cellular processes like cell survival, cell metabolism and immune responses. Studies suggest that viruses utilize host Akt signaling for their successful replication. Viral protein activates Akt signaling to inhibit proapoptotic factors by activating transcription factors such as FoxO1 and promotes cell survival for their better propagation (1). Virus also employs PI3K-Akt signaling to inhibit interferon response (2). Literature also suggests that Akt phosphorylation helps with assembly of RNA-dependent RNA polymerase replication complex of the viruses (3). RNA viruses, like severe acute respiratory syndrome coronavirus-2 (SARS CoV-2), are also reported to employ Akt signaling for their propagation as well (4). Akt inhibitors significantly inhibit replication and propagation of SARS CoV-2 via Akt-mTOR-HIF1α axis (5). Inflammatory mediators such as high-mobility group box 1 (HMGB1) protein elevate pAkt and in turn increase expression of angiotensin-converting enzyme 2 (ACE2) (6), the receptor for SARS CoV-2 (7). Akt regulates the expression of ACE2 as well as transmembrane serine protease 2 (TMPRSS2), mediators of SARS CoV-2 invasion mechanism (8). In contrast, the host also modulates Akt signaling to regulate immune responses. Akt phosphorylation potentiates translation of interferon-stimulated genes (ISGs) through actions of mTOR and p70S6K, which stimulate ISG translation (2). Akt also plays a role in stimulating secretion of inflammatory cytokines by activating NFkB (9). Akt signaling leads to activation of host innate immune responses including secretion of inflammatory cytokines. Akt activation results in increased production of pro-inflammatory cytokines like IL6 and IL1β in virus infected cells (10). Influenza infected cells produce IL8 and RANTES in a PI3K/Akt-dependent manner, (11) and viral activation of Akt potentiates the expression of interferon-β (12). Besides, hypoxia inducible factor (HIF)1α, which is elevated in hypoxic microenvironment, is also known to promote the viral infection. SARS-CoV-2 ORF3a protein induces HIF1α function, which subsequently facilitates viral infection and triggers cytokine production in COVID-19 patients (13). HIF1α is also an important activator of glycolysis and inflammatory responses, implying the role of HIF1α in the pathogenesis of COVID-19 (14).

In our recent works, we described that α-ketoglutarate (αKG), a co-factor of prolyl hydroxylase 2 (PHD2), inhibited both phospho-Akt (pAkt) (15) and HIFα (16) by augmenting the prolyl hydroxylation activity of PHD2. αKG significantly inhibited both pAkt and HIFα to abrogate pro-inflammatory response of leukocytes as well as pro-thrombotic role of platelets in mice (15). Also, we described that dietary αKG reduces lung inflammation in SARS CoV-2 infected hamsters (15). However, our above study was limited in describing the anti-inflammatory effects of αKG. In this study, we describe a potential anti-viral role of αKG in inhibiting viral replication, and its rescue effect on inflammation and thrombosis in lungs as well as in circulation of SARS CoV-2 infected animals, without affecting the anti-viral response of the IgG as well as IFNγ+CD4+ T lymphocytes. Dietary supplementation of αKG also restored the O_2_ saturation percentage in circulation of SARS CoV-2 infected animals, suggesting safe and potent usage of this metabolite.

## RESULTS

### αKG supplementation inhibits SARS CoV-2 replication in a concentration dependent manner in parallel with the inhibition of pAkt in cells *in vitro*

α-Ketoglutarate (αKG) inhibits Akt phosphorylation (pAkt) (15) and HIFα function (16) by augmenting the PHD2-mediated hydroxylation of the proline resides of these substrates. Here we show that octyl αKG (cell membrane permeable form of αKG) significantly inhibits SARS CoV-2 replication and infection (negative strand of viral RNA was estimated by RT-PCR and calculated as Log10 fold change, p<0.01, Fig. 1A; and viral spike protein was measured using microscopy and expressed as mean fluorescence intensity, MFI, p<0.0001, Fig. 1B-C) in Vero E6 cells at 24 hr of infection. Upon infection with 0.01 MOI of virus, the SARS CoV-2 replication in these cells was found to be highest at 24 hr of infection, compared to other time points such as 12 and 36 hr (Fig. S1A). Therefore, subsequent experiments were performed at 24 hr. A known Akt inhibitor, triciribine (TCN) also inhibited the viral replication (Log10 fold change of viral genome, p<0.05, Fig. 1A; spike protein expression in MFI, p<0.0001, Fig. 1B-C). αKG decreased the pAkt in these cells in a concentration dependent manner (western blot densitometry fold change of pAkt-Ser473, p<0.0001, pAkt-Thr308, p<0.05, Fig. 1D), parallelly with the decrease in viral replication (Fig. 1A). αKG also decreased expression of other substrates of the PHD2 such as hypoxia inducible factor 1α (HIF1α) and HIF2α (densitometry fold change, p<0.05 and p<0.01, Fig. 1D). Further, we describe that both αKG and TCN decreased ACE2 expression in these cells (densitometry fold change, for αKG or TCN, p<0.05, Fig. 1D), suggesting the role of pAkt in ACE2 expression and SARS CoV-2 entry into the cells. An experiment was performed in human monocytic U937 cell line, transiently expressing adenovirus+hACE2 (Fig. 2A). αKG treatment inhibited the viral replication (Log10 fold change of viral genome, p<0.05, Fig. 2A, C) and spike protein (MFI, p<0.0001, Fig. 2E-F) of SARS CoV-2, in parallel with a decreased pAkt-Ser473 and pAkt-Thr308 (densitometry fold change, p<0.05 and p<0.01, Fig. 2I) in infected-U937 cells. Since αKG works through PHD2 axis, we performed the above experiment in PHD2-knockdown (KD) U937 cell, transiently expressing hACE2 (Fig. 2B). As expected, we did not observe a decrease in αKG-mediated inhibition of SARS CoV-2 replication (viral genome, Fig. 2D, and spike protein, Fig. 2G-H) in parallel with an unaltered pAkt in PHD2-KD cells (Fig. 2J). Whereas, TCN inhibited the viral replication and propagation (Log10 fold change of viral genome, p<0.05, and spike protein MFI, p<0.0001, Fig. 2G-H) in parallel with the downmodulation of pAkt-Ser473 and pAkt-Thr308 (densitometry fold change, p<0.05 and p<0.05, Fig. 2J) in PHD2-KD cells. Above observations establish that αKG works through PHD2 axis to inhibit pAkt.

**Fig. 1:**
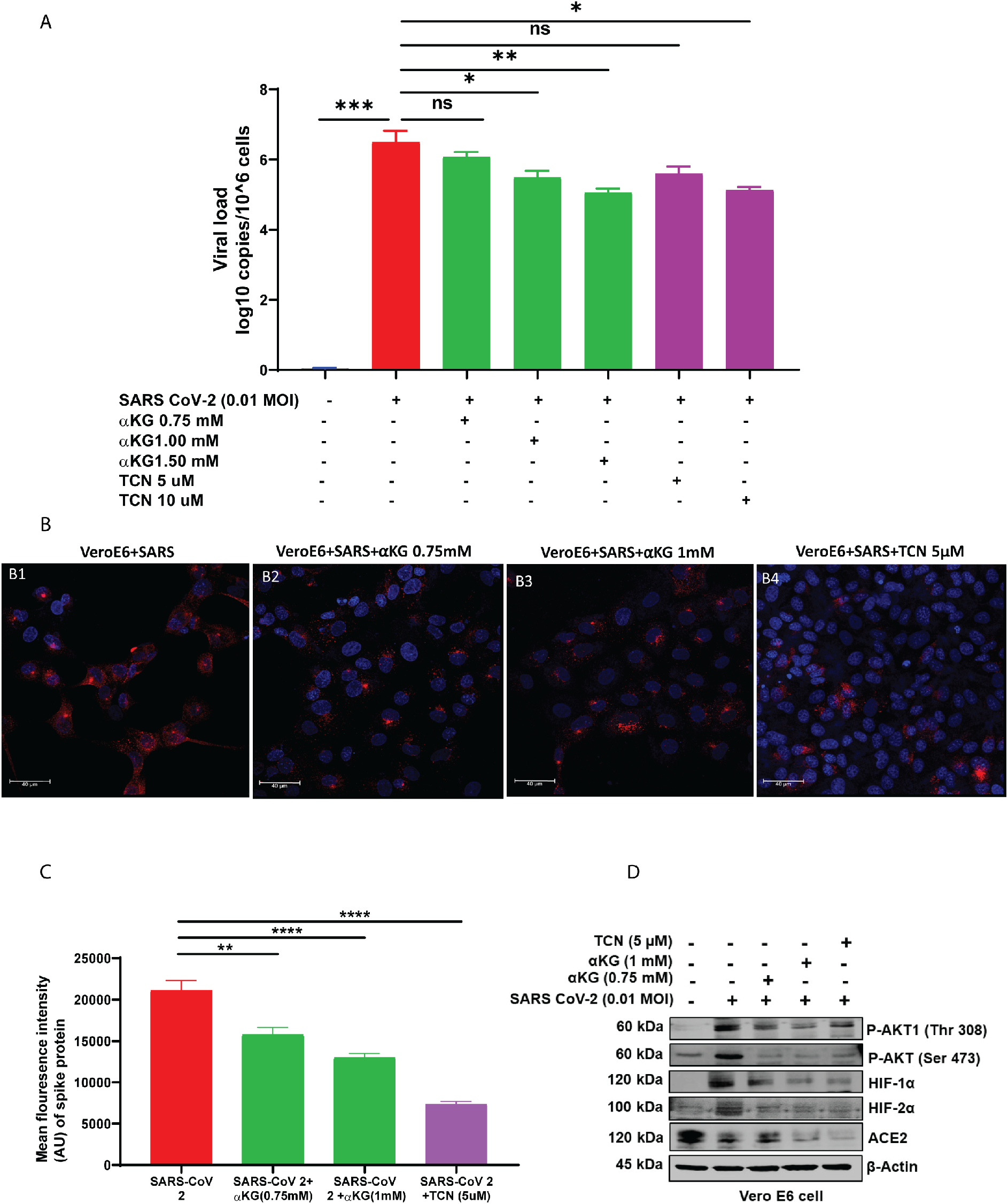
Viral replication in Vero E6 cells is decreased by αKG supplementation: (A) Vero E6 cells were infected with SARS CoV-2 (0.01 MOI) for 24 hr in presence of αKG or TCN. αKG decreased viral genome negative strand (measured by RT-PCR) dose-dependently. Data are from three independent experiments and represented as mean ± SEM (Kruskal-Wallis test followed by Dunn’s multiple comparison post-test), *P<0.05, **P<0.01, ***P<0.001, and ns=non-significant. (B) Spike protein expression (red puncta) was measured in above mentioned cells using confocal microscopy. (C) MFI of Spike protein is quantified from 50 cells from different experiments. Values are represented as mean ± SEM (Kruskal-Wallis test followed by Sidak’s multiple comparison post-test), **P<0.01, ****P<0.0001. (D) Protein expression of pAkt1-Thr308, pAkt-Ser473, HIF1α, HIF2α and ACE2 was detected by western blot. The expression of all above proteins were decreased upon treatment with either αKG or TCN in SARS CoV-2 infected Vero E6 cells. Densitometry quantification from 3 different experiments is described in Fig. S2A-E (Kruskal-Wallis test followed by Dunn’s multiple comparison post-test), *P<0.05, **P<0.01, ***P<0.001, ****P<0.0001 and ns=non-significant.

**Fig. 2:**
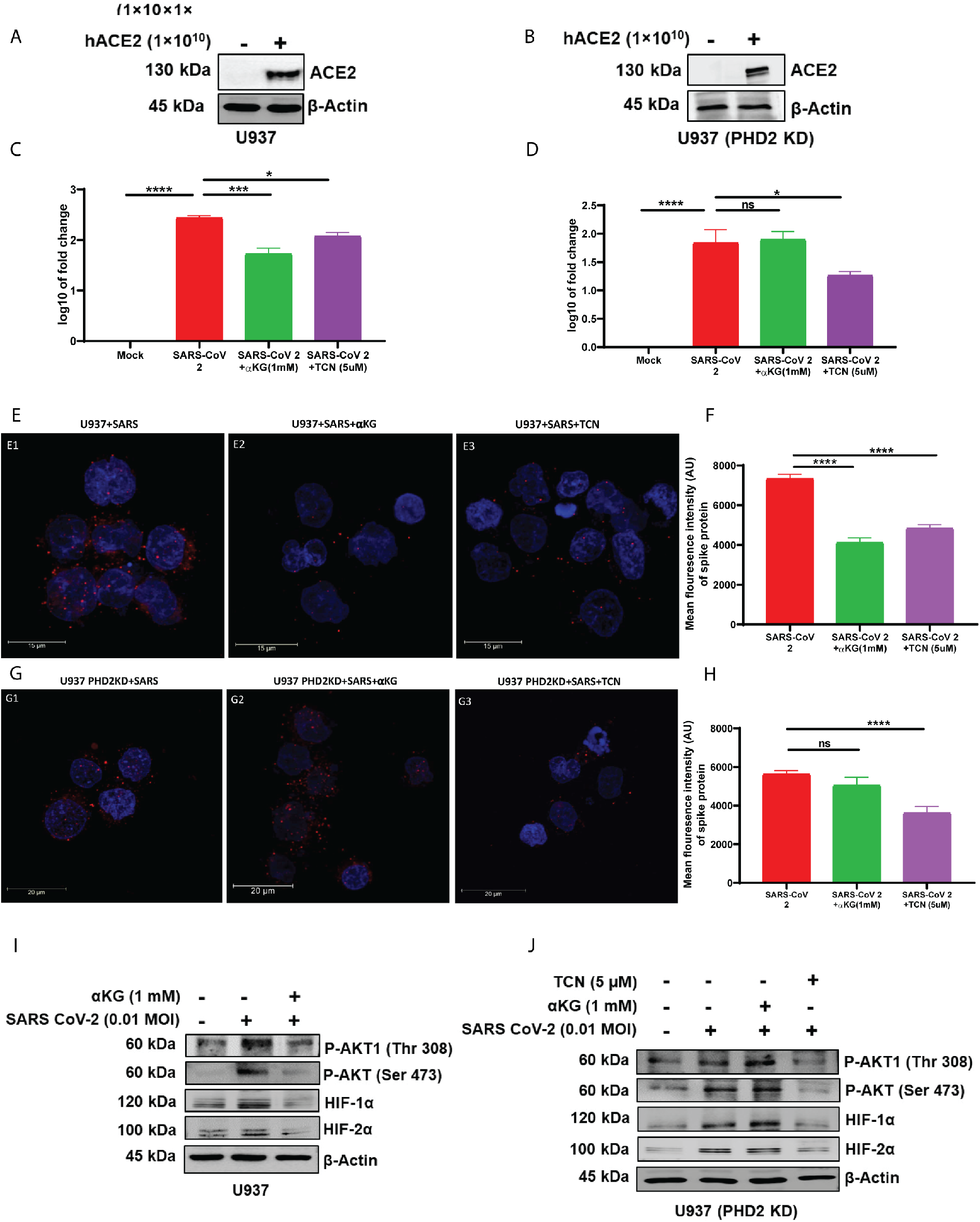
αKG inhibits pAkt via PHD2 axis: (A-B) ACE2 was expressed transiently in (A) monocytic U937 cell line or (B) PHD2-knockdown U937 cells using a human ACE2 adenovirus construct. (C, E-F) αKG or TCN decreased SARS CoV-2 viral replication [negative strand of viral RNA was measured and viral spike protein (red puncta) was assessed] in wildtype U937 cells. (D, G-H) αKG was unable to decrease viral replication (negative strand of viral RNA and Spike protein) in PHD2-knockdown U937 cells. TCN decreased viral replication in PHD2-KD U937 cells. Data are from three independent experiments (2C-D) and represented as mean ± SEM (one-way ANOVA using Kruskal-Wallis’s post-test), *P<0.05, ***P<0.001, ****P<0.0001 and ns=non-significant. (E-H) MFI of Spike protein was quantified from 50 cells from different experiments. Values are represented as mean ± SEM (one-way ANOVA using Kruskal-Wallis’s post-test), ****P<0.0001 and ns=non-significant. (I-J) Expression of pAkt1-Thr308, pAkt-Ser473, HIF1α and HIF2α was decreased upon treatment with αKG in SARS CoV-2 infected wild-type U937 cells, but found unaltered in PHD2-KD U937 cells. Densitometry quantification from 3 different experiments is described in Fig. S2H-O (Kruskal-Wallis test followed by Dunn’s multiple comparison post-test), *P<0.05, **P<0.01, ***P<0.001, ****P<0.0001 and ns=non-significant.

### Dietary αKG supplementation inhibits SARS CoV-2 propagation in hamster in parallel with the inhibition of pAkt

SARS CoV-2 infected hamsters were fed with human dietary grade αKG (40 mg/100 g body wt./day), via oral gavage, from the next day after nasal infection of SARS CoV-2 till 4 days post infection (dpi), and experiment was terminated either 5 or 9 dpi, described in schematic Fig. 3A). αKG significantly inhibited viral load (fold change of genome copy, p<0.01 and p<0.05, at 5 and 9 dpi accordingly, Fig. 3B-C) and spike protein expression (immunostaining intensity, p<0.0001 at 5 dpi, Fig. 3D) in the lungs of the hamsters. No difference was observed in the expression of spike protein in the lungs of the animals from infected vs. infected+αKG group at 9 dpi (Fig. 3F-G). Body wt. was decreased following viral infection, although, no significant rescue in wt. loss was observed in αKG group (Fig. S6). A significant downmodulation of pAkt-Thr308 and pAkt-Ser473 (WB densitometry fold change, p<0.05 and p<0.0001, Fig 3H, and immunostaining intensity of pAkt, p<0.01, Fig. 3I-J) was observed in infected hamsters compared to infected+αKG group at 5 dpi. αKG also inhibited the HIF2α expression (WB densitometry fold change, p<0.001, Fig. 3H; immunostaining intensity, p<0.0001, Fig. 3K-L) in the lungs of the infected animals after αKG supplementation.

**Fig 3:**
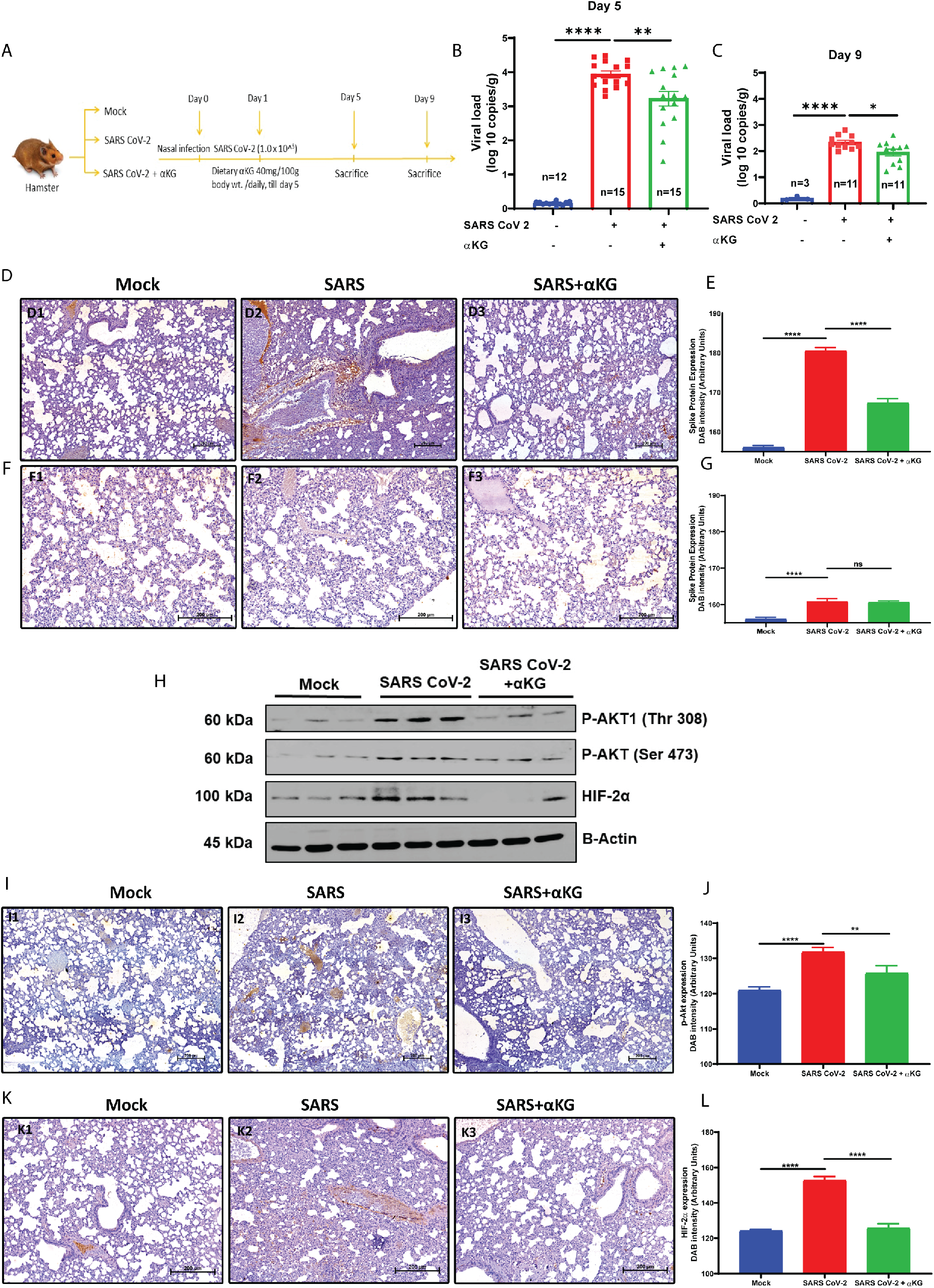
αKG rescues hamsters from SARS CoV-2 infection: (A) Schematic representation of experimental protocol. Hamsters were given SARS CoV-2 infection via nasal route and αKG was supplemented from 1 dpi. Hamsters were sacrificed at 5 dpi and 9 dpi and following parameters were measured. (B-C) αKG reduces the SARS CoV-2 viral replication in hamster lung (measured by RT-PCR, 5 dpi and 9 dpi). Data are from 12 mock and 15 each infected and infected+αKG at 5 dpi, and 3 mock and 11 each infected and infected+αKG at 9 dpi. Each dot represents individual value. Data are represented as mean ± SEM (one-way ANOVA using Sidak’s post-test), **P<0.01 and ****P<0.0001. (D-G) Expression of Spike protein in hamster lungs was reduced upon αKG supplementation at 5 dpi (3D) and 9 dpi (3F) measured by IHC. DAB intensity was quantified at 5 dpi (3E) and 9 dpi (3G). Data are from 10 fields from different animals. Values are represented as mean ± SEM (one-way ANOVA using Bonferroni’s post-test), ****P<0.0001 and ns=non-significant. (H) Expression of pAkt1-Thr308, pAkt-Ser473 and HIF2α was measured in SARS CoV-2 infected hamsters supplemented with αKG. Densitometry quantification from different animals is described in Fig. S2P-R. (Kruskal-Wallis test followed by Dunn’s multiple comparison post-test), *P<0.05, **P<0.01, ***P<0.001, ****P<0.0001, and ns=non-significant. (I) pAkt expression in αKG treated SARS CoV-2 infected lung measured using IHC. (K) HIF2α expression in αKG treated SARS CoV-2 infected lung measured using IHC at 5 dpi. (J, L) DAB intensity quantification of pAkt and HIF2α expression. Data are from 10 fields from different animals. Values are represented as mean + SEM (one-way ANOVA using Bonferroni’s post-test), **P<0.01 and ****P<0.0001.

### Dietary αKG rescues exaggerated inflammation and thrombus formation in the lungs of SARS CoV-2 infected hamsters

Dietary supplementation of αKG significantly decreased SARS CoV-2 induced accumulation of inflammatory cells at 5 and 9 dpi in the alveolar spaces (percentage cellularity, p<0.0001 and p<0.0001 at 5 and 9 dpi, Fig. 4A-B and Fig. 4C-D) and thrombus formation in the micro blood vessels (thrombus score arbitrary unit, p<0.0001 and p<0.05 at 5 and 9 dpi, Fig. 4E-F and Fig. 4G-H). The accumulation of inflammatory cells and microthrombi are the known causes of acute respiratory distress syndrome (ARDS) in COVID-19.

**Fig 4:**
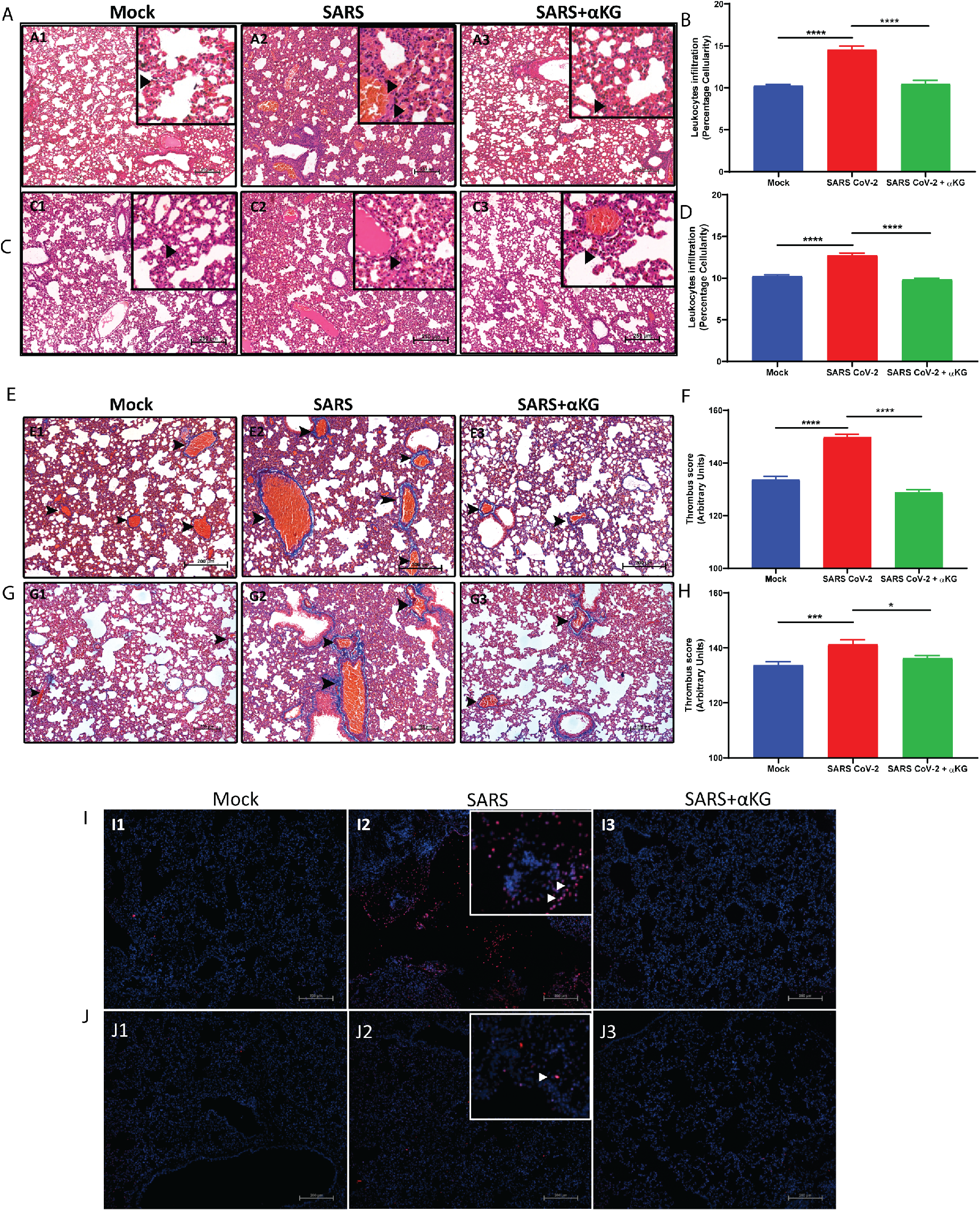
αKG rescues inflamed lungs in SARS CoV-2 infected hamsters: (A-D) H&E staining of lung was used to assess leukocyte accumulation at 5 dpi (4A) and 9 dpi (4C). Score was calculated as percentage cellularity at 5 dpi (4B) and 9dpi (4D), arrows indicate cell accumulation. Scale bar 250 mm. αKG supplementation reduced inflammation in infected hamsters. Data are from 10 fields from different animals. Values are represented as mean ± SEM (one-way ANOVA, using Bonferroni’s post-test **** P<0.0001). (E-F) MT staining of lung was used for assessing clot formation at 5 dpi (4E) and 9 dpi (4G), arrows indicate thrombus or clot. Score was calculated using color deconvolution protocol in image-J software at 5 dpi (4F) and 9 dpi (4H). Scale bar 200 mm. αKG treatment reduced thrombosis in infected hamsters. Data are from 10 fields from different animals. Values are represented as mean ± SEM (one-way ANOVA, using Bonferroni’s post-test *P<0.05, ***P<0.001 and **** P<0.0001). (I-J) TUNEL assay in the lungs of infected hamsters was used to assess apoptotic cells (red puncta). αKG treatment reduced apoptosis in the lungs of infected hamsters at 5 dpi (Fig. 4I) and 9 dpi (Fig. 4J). More images of Fig. 4I-J are described in Fig. S5.

### Dietary αKG rescues lungs from apoptotic tissue damage in SARS CoV-2 infected hamsters

SARS CoV-2 infection induced the apoptotic tissue damage in lungs of SARS CoV-2 infected animals 5 dpi (Fig. 4I) and 9 dpi (Fig. 4J). Almost no apoptotic tissue damage was observed in infected hamsters after dietary αKG supplementation, suggesting a rescue effect of the metabolite.

### Dietary αKG does not interfere with anti SARS CoV-2 antibody generation and percentage of the interferon-γ positive T cells in infected hamsters

Our above observations describe the anti-thrombotic and anti-inflammatory effects as well as anti-apoptotic role of dietary supplementation of αKG in SARS CoV-2 infected hamsters. We thereafter investigated the effect of αKG on adaptive immune parameters in these animals, if any. We observed similar plasma levels of IgG against SARS CoV-2 RBD protein 5 dpi (Fig. 5A-B) and 9 dpi (Fig. 5C-D) between infected and infected+αKG groups. We observed a similar efficacy of the plasma containing antibody from αKG supplemented animals in neutralizing the SARS CoV-2, compared to only virus infected group (Fig. 5E-F), suggesting no interference of αKG on antibody response. We investigated the T lymphocyte profile between groups. Dietary αKG did not alter the already elevated percentage of interferon-γ positive (IFNγ+) CD4+ (Fig. 5G-H) and IFNγ+CD8+ (Fig. 5I-J) T cells in the spleen compared to only virus infected group 5 and 9 dpi respectively, thus suggesting a safe usage of the metabolite without affecting antibody response, specifically neutralizing ability against SARS CoV-2. No significant difference was observed in platelet, RBC and WBC counts in blood between infected and infected+αKG groups (Fig. S4).

**Fig. 5:**
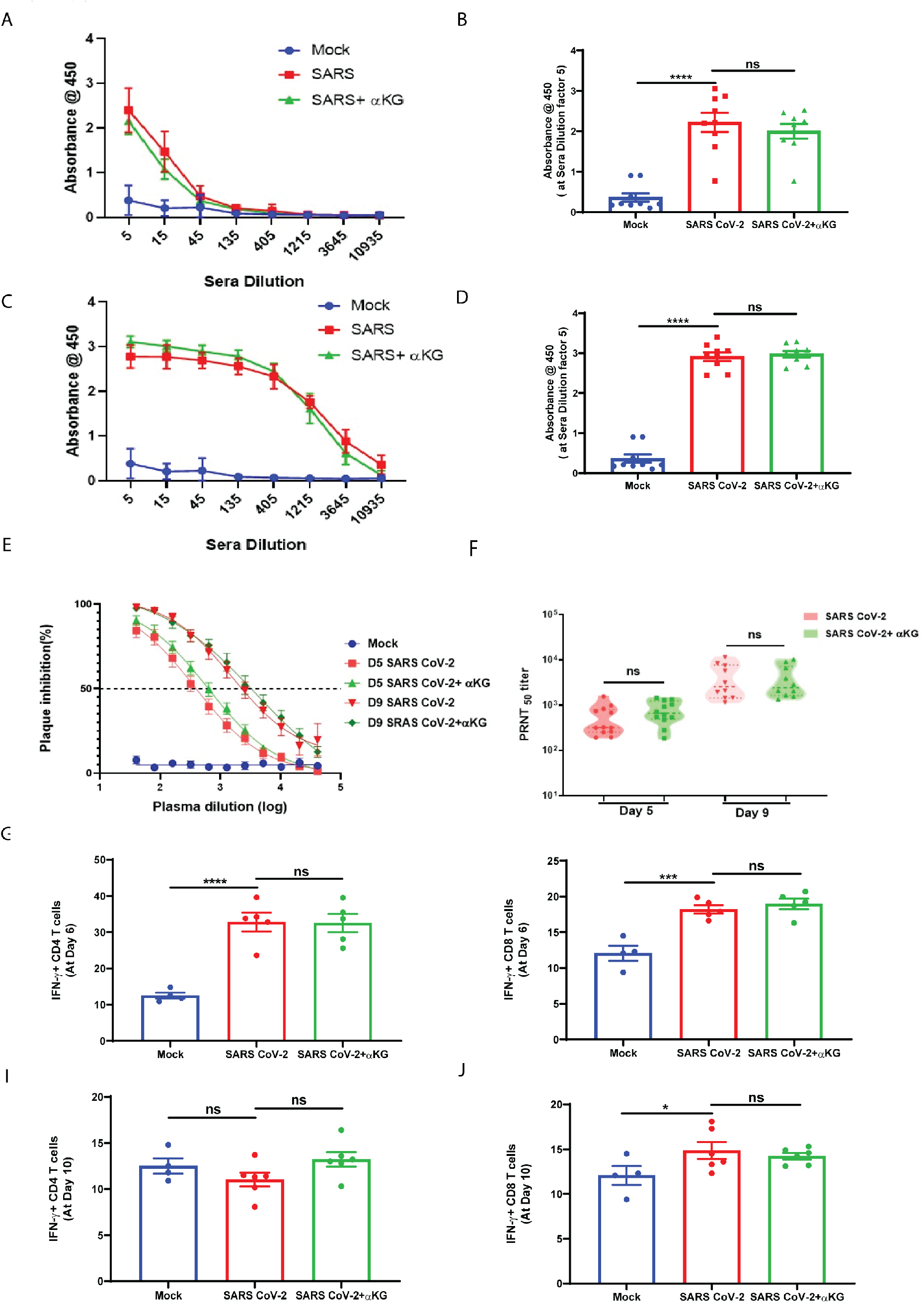
αKG does not interfere with anti SARS CoV-2 antibody generation and percentage of the interferon-γ positive T cells in infected hamsters. (A-D) Anti-SARS RBD antibody quantification at increasing sera dilution showed no difference between infected and infected+αKG group at 5 dpi (5A) and 9 dpi (5C). Absorbance at sera dilution 5 at 5 dpi (5B) and 9 dpi (5D). Data are from 9 animals in each group. Values are represented as mean ± SEM (one-way ANOVA, using Bonferroni’s post-test, **** P<0.0001 and ns=non-significant). (E-F) Neutralization antibody was measured at increasing sera dilution using PRNT50 assay showing no difference between above groups at 5 dpi and 9 dpi. Data are from 12 animals in each group at 5 dpi and 10 animals in each group at 9 dpi. Values are represented as mean ± SEM (one-way ANOVA, ns=non-significant). (G, I) Flow cytometry analysis of IFNγ+ CD4 T cells showing no difference in the αKG treated hamster spleen as compared to only SARS CoV-2 infected hamster spleen at 5 dpi (5G) and 9 dpi (5I). Gating strategy is mentioned in fig.S7. Data are from 4 animals in mock and 5 animals each in SARS CoV-2 infected and αKG treated animals at 5 dpi and 6 animals each in SARS CoV-2 infected and αKG treated animals at 9 dpi. Values are represented as mean ± SEM (one-way ANOVA, using Bonferroni’s posttest, ****P<0.0001 and ns=non-significant). (H-I) Flow cytometry analysis of IFNγ+ CD8 T cells showing no difference in the αKG treated hamster spleen as compared to only SARS CoV-2 infected hamster spleen at 5 dpi (5G) and 9 dpi (5I). Data are from 4 animals in mock and 5 animals each in SARS CoV-2 infected and αKG treated animals at 5 dpi and 6 animals each in SARS CoV-2 infected and αKG treated animals at 9 dpi. Values are represented as mean ± SEM (one-way ANOVA, using Bonferroni’s post-test, *P<0.05, ***P<0.001 and ns=non-significant).

### Dietary αKG inhibits exaggerated inflammation and pro-thrombotic events in the lungs of the mice infected with SARS CoV-2

A similar study was performed in balb/c mice, transiently expressing hACE2 in the upper respiratory tract (by intranasal treatment of hACE2 adenovirus). The mice were infected with SARS CoV-2 and fed with αKG (8 mg/20 g body wt./day, via oral gavage, from the next day after nasal infection of SARS CoV-2 till 4 dpi, and experiment was terminated either 5 and 9 dpi, described in schematic Fig. 6A). A significant inhibitory effect of αKG on viral load (PCR quantification) was observed 5 dpi (fold change, p<0.001, Fig. 6B). The infection was reduced 9 dpi and no difference was observed between infected vs infected+αKG group (Fig. 6C).

αKG supplementation reduced the accumulation of inflammatory cells 5 and 9 dpi in the alveolar spaces (percentage cellularity, p<0.0001 for both 5 and 9 dpi, Fig. 6D-E and Fig. 6H-I) and thrombus formation in the micro blood vessels (thrombus score arbitrary unit, p<0.0001 and p<0.0001 at 5 dpi, Fig. 6F-G, p=ns at 9 dpi, Fig. 6J-K) in the lungs of infected animals. αKG supplementation significantly reduced the elevated counts of leukocytes (percentage, p<0.001, Fig. 6L, and p<0.0001, Fig. 6N) and leukocyte-platelet aggregates (percentage, p<0.01, Fig. 6M, and p<0001, Fig. 6O) in the lungs of infected mice at 5 and 9 dpi.

**Fig. 6:**
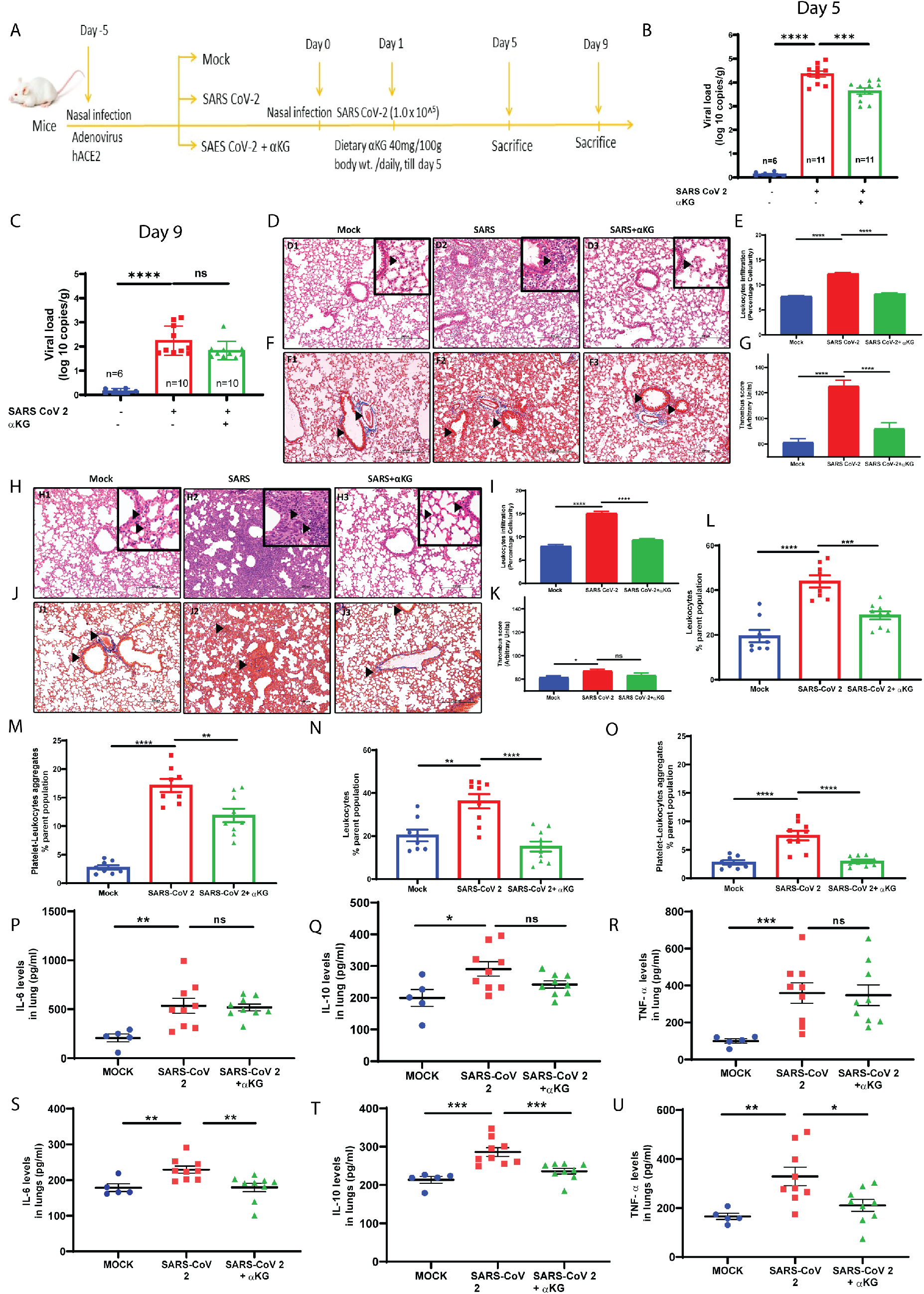
αKG inhibits inflammation and thrombosis in the lungs of SARS CoV-2 infected mice. (A) Schematic representation of experimental protocol. Balb/c mice were given human ACE2 adenovirus construct via intra nasal route for transient expression of ACE2, after 5 days mice were given SARS CoV-2 infection via nasal route and αKG was supplemented from 1 dpi. Mice were sacrificed at 5 dpi and 9 dpi and following parameters were measured. (B-C) αKG reduces the SARS CoV-2 viral replication in mice (measured by RT-PCR, 5 dpi and 9 dpi) Data are from 6 mock and 11 each from infected and infected+αKG group at 5 dpi and from 6 mock and 10 each from infected and infected+αKG group at 9 dpi. Values are represented as mean ± SEM (one-way ANOVA using Sidak’s post-test), ***P<0.001, ****P<0.0001 and ns=non-significant. (D-E, H-I) H&E staining of lung was used for assessing leukocyte accumulation 5 dpi (6D) and 9 dpi (6H). Score was calculated as percentage cellularity at 5 dpi (6E) and 9 dpi (6I), showing a reduced inflammation by αKG treatment. Data are from 10 fields from different animals. Values are represented as mean ± SEM (oneway ANOVA, using Bonferroni’s post-test **** P<0.0001). (F-G, J-K) MT staining of lung was used to assess thrombus score at 5 dpi (6F) and 9 dpi (6J). Score was calculated using color deconvolution protocol in image-J software at 5 dpi (6G) and 9 dpi (6K), showing reduced thrombosis in αKG-treated hamsters. Data are from 10 fields from different animals. Values are represented as mean ± SEM (one-way ANOVA, using Bonferroni’s post-test *P<0.05, **** P<0.0001 and ns=non-significant). (L-O) Leukocytes population and leukocyte-platelet aggregation population in single cell suspension of lung was assessed using flow cytometry. αKG treatment reduced leukocytes number in lungs at 5 dpi (6L) and 9 dpi (6N). αKG treatment also reduced leukocytes-platelet aggregates in lungs at 5 dpi (6M) and 9 dpi (6O). Data are from 8 animals in mock and 8-10 animals each in SARS CoV-2 infected and αKG-treated. Values are represented as mean ± SEM (one-way ANOVA, using Bonferroni’s post-test **P<0.01, ***P<0.001, **** P<0.0001). (P-U) IL6, IL10 and TNFα were measured in lungs of SARS CoV-2 infected animals and αKG-treated animals at 5 dpi (6P-6R) and 9 dpi (6S-6U). Data are from 6 animals in mock and 9 animals each in infected and infected+αKG group. Values are represented as mean ± SEM (one-way ANOVA, using Bonferroni’s post-test *P<0.05, **P<0.01, ***P<0.001 and ns=non-significant).

αKG supplementation significantly rescued the elevated levels of cytokines such as IL6, IL10 and TNFα (concentration in pg/ml, p<0.01, p< 0.001 and p<0.05, Fig. 6S-U) 9 dpi in the lungs of infected mice. The rescue effect of αKG was not prominent for the above cytokines 5 dpi(Fig. 6P-R).

### Dietary αKG inhibits elevated cytokines and platelet microparticles in circulation in the mice infected with SARS CoV-2

αKG supplementation significantly reduced the elevated levels of cytokines, IL6, IL10 and TNFα in plasma 5 dpi (concentration in pg/ml, p<0.05, p< 0.01 and p<0.01, Fig. 7A-C) as well as 9 dpi (p<0.01, p< 0.001 and p<0.01, Fig. 7D-F) of SARS CoV-2 infected mice. αKG supplementation also reduced the platelet (CD41a+) microparticles (thrombogenic as well as platelet activation marker) 5 dpi (calculated as percentage event, p<0.001, Fig. 7G) as well as 9 dpi (p<0.001, Fig. 7H), suggesting a potent anti-inflammatory and anti-thrombotic role of the metabolite.

**Fig. 7:**
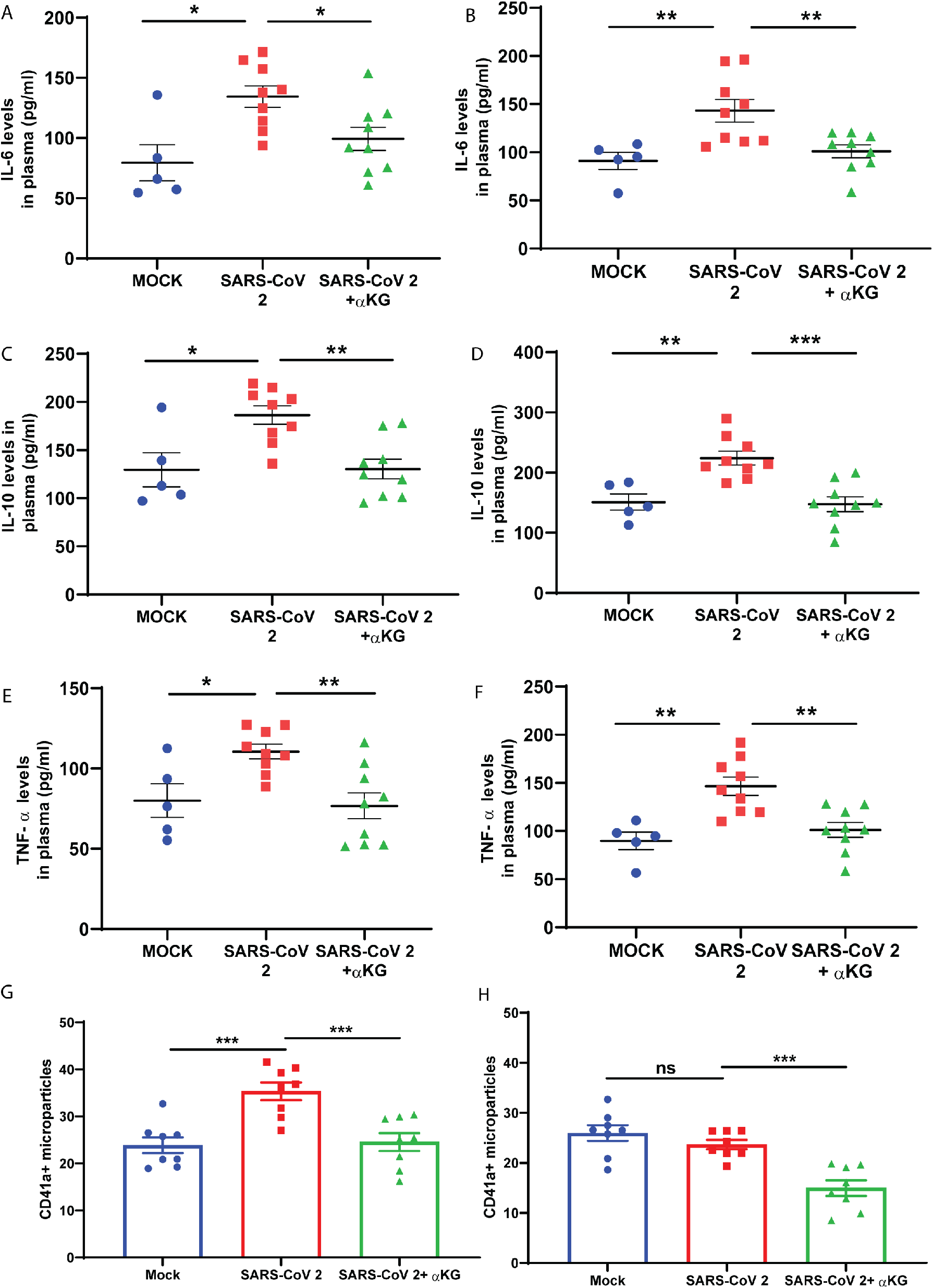
αKG inhibits elevated cytokines and platelet microparticles in SARS CoV-2 infected mice. (A-F) IL6, IL10 and TNFα were measured in plasma of infected and infected+αKG group at 5 dpi (7A-C) and 9 dpi (7D-F). αKG reduced above cytokines levels in plasma. Data are from 6 animals in mock and 9 animals each in infected and infected+αKG group. Values are represented as mean ± SEM (one-way ANOVA, using Bonferroni’s posttest *P<0.05, **P<0.01, ***P<0.001). (G-H) CD41a+ microparticles were measured using flow cytometry in plasma of SARS CoV-2 infected animals and αKG-treated animals at 5 dpi (7G) and 9 dpi (7H). αKG reduced CD41a+ microparticles in the plasma. Data are from 8 animals in each group. Values are represented as mean ± SEM (one-way ANOVA, using Bonferroni’s post-test, **P<0.01, ***P<0.001 and ns= non-significant).

### Dietary αKG restores SpO_2_ percentage in circulation of hamsters and mice infected with SARS CoV-2

SARS CoV-2 infection causes clinical symptoms like hypoxemia resulting in a decrease in the oxygen pressure saturation (SpO_2_) in blood. We measured the percentage SpO_2_ in the above infected animals in awake condition. SARS CoV-2 infected hamsters had a decreased SpO_2_ percentage in most of the animals (7 or 8 out of 10) between 2 dpi and 7 dpi. Interestingly, the αKG supplementation maintained normal SpO_2_ in the infected animals (percentage SpO_2_ at 4 dpi, p<0.05, Fig. 8A). Unlike the hamsters, the infected balb/c mice did not show a gradual decrease in SpO_2_ except 5 dpi. The αKG supplementation maintained the normal SpO2 in mice (percentage SpO_2_ at 5 dpi, p<0.01, Fig. 8B). Thus, αKG supplementation rescued the animals from decrease in oxygen percentage in the blood following SARS CoV-2 infection.

**Fig. 8:**
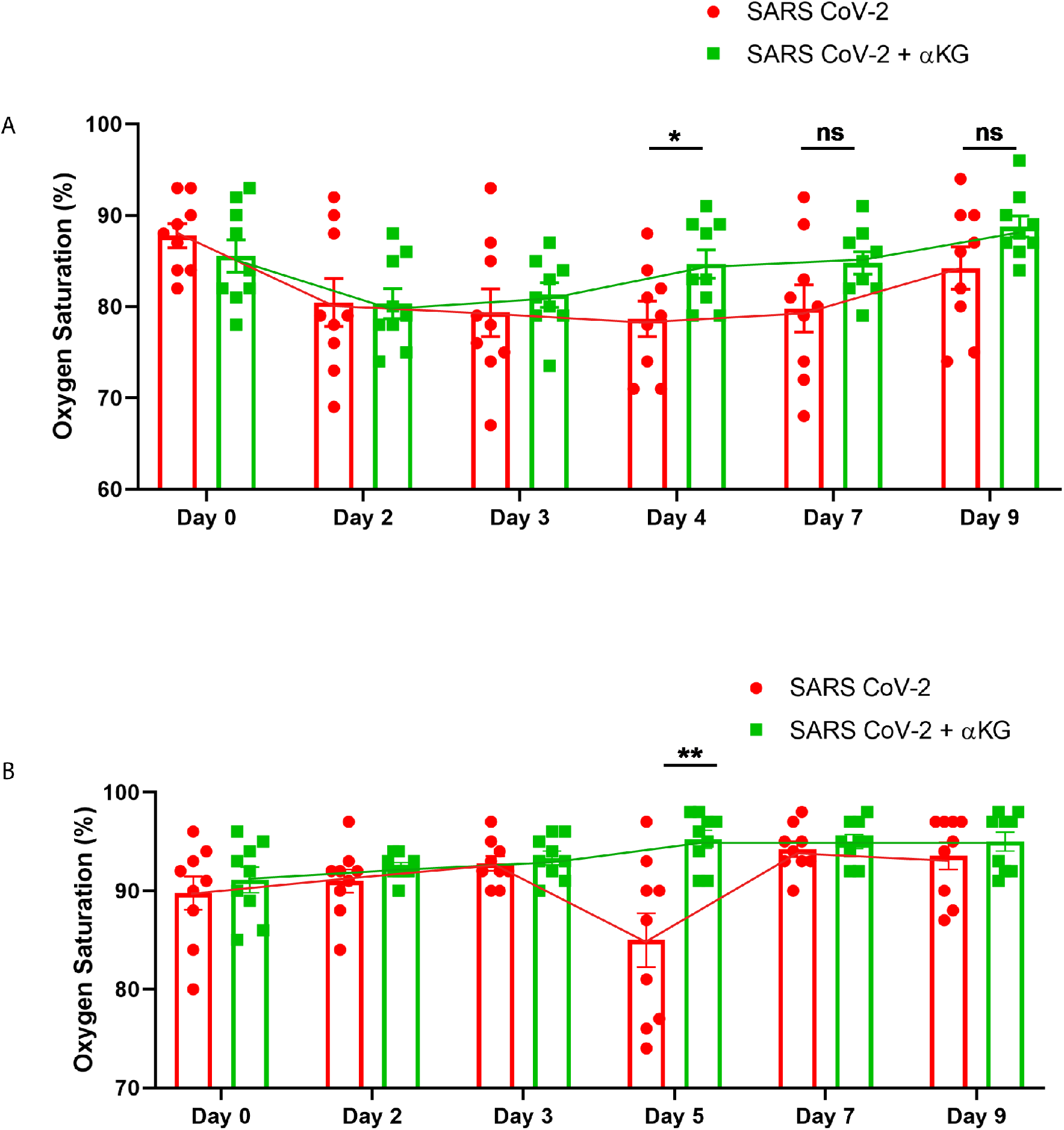
αKG restores SpO2 percentage in circulation of the hamsters and mice infected with SARS CoV-2. (A-B) Oxygen saturation was measured in circulation of the infected vs infected+αKG hamsters (8A) and balb/c mice (8B) using pulse oximeter. Oxygen saturation was dropped during 2 dpi to 9 dpi in infected animals. But αKG rescued it. Data are from 9 animals in each group. Values are represented as mean ± SEM (student’s t test, *P<0.05 and ns=non-significant).

## DISCUSSION

In this study, we describe that 1) phosphorylation of Akt in host cells is directly correlated with replication of SARS CoV-2. 2) αKG supplementation inhibits SARS CoV-2 replication by PHD2-mediated inactivation of pAkt. 3) Dietary supplementation of αKG, on one hand, inhibited viral replication, while on the other hand, reduced inflammation, thrombosis and apoptotic cell death in lungs of SARS CoV-2 animals without affecting the interferon-γ positive (IFNγ+) CD4 and CD8 T-cell profile and antibody response. 4) αKG supplementation restored normal oxygen saturation in infected animals (schematic Fig. 9).

**Fig. 9:**
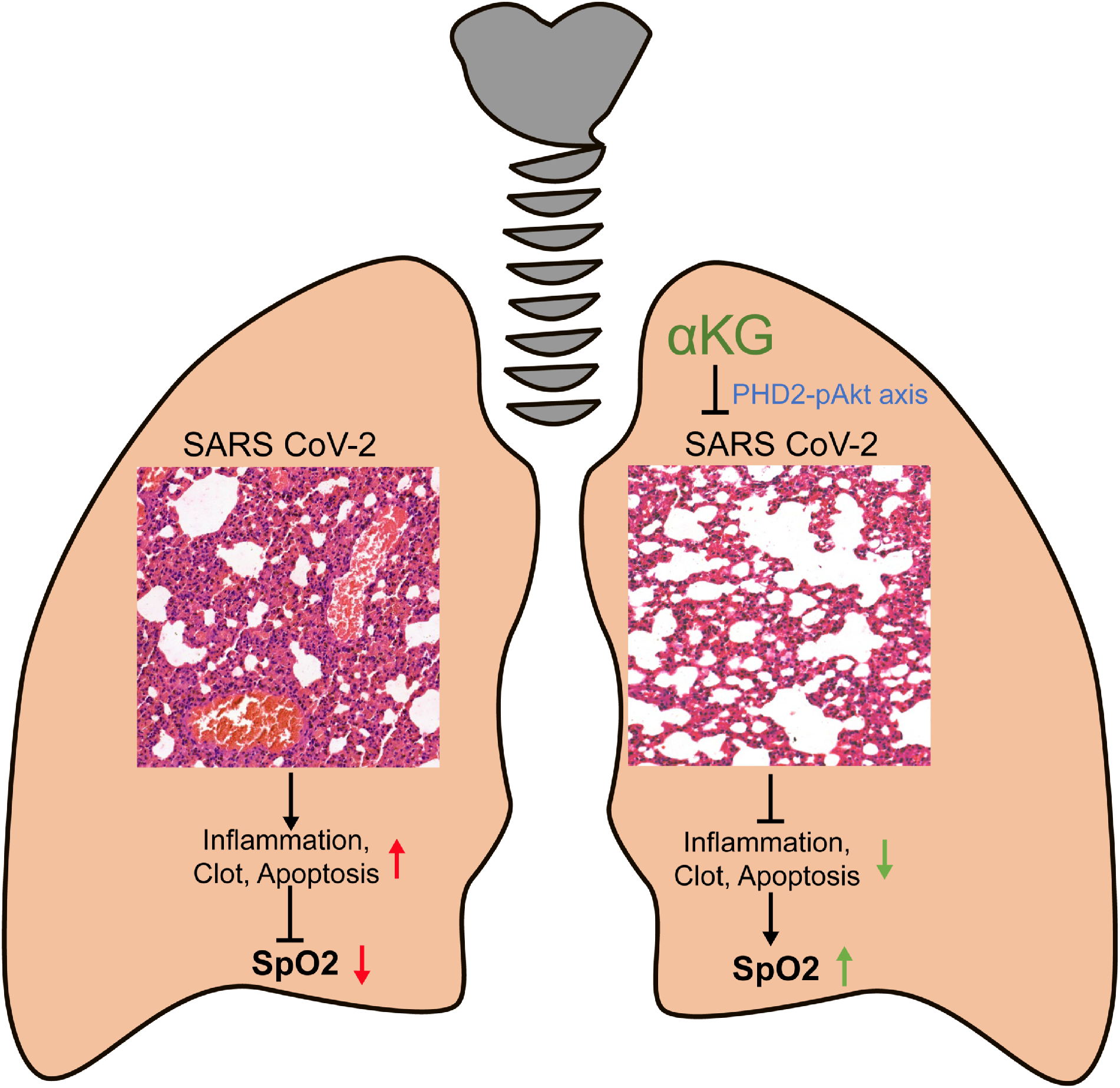
**Schematic** describes an elevated inflammation, thrombosis and apoptotic tissue damage in the lungs of SARS CoV-2 infected animals leading to a condition called hypoxemia or less O_2_ in the circulation. In contrast, αKG supplementation rescues animals from SARS CoV-2 mediated above clinical events by downmodulating PHD2-mediated Akt phosphorylation, resulting in a restoration of normal oxygen pressure saturation (SpO_2_) in blood.

Viruses utilize host Akt signaling for successful replication. Viral protein activates Akt signaling thus inhibiting proapoptotic factors and promoting host cell survival for better viral propagation (17). SARS-CoV N protein activates Akt phosphorylation to promote viral replication in Vero E6 cells (18). Our data also describe a direct correlation between elevated Akt phosphorylation (pAkt) and increased replication of SARS CoV-2 in Vero E6 cells. Importantly, the supplementation of αKG, a metabolite of Krebs cycle, inhibited the SARS CoV-2 replication in a concentration dependent manner. In a recent work, we have described that αKG supplementation augments the prolyl hydroxylation activity of PHD2 and degrades proline residues of pAkt, and in turn inactivating pAkt (15). Therefore, the αKG mediated inhibition of pAkt activity may be a plausible reason for inhibition of SARS CoV-2 replication. Several other studies have described the direct association of PI3K/Akt signaling of host cells with SARS CoV-2 replication. The Akt inhibitor like MK-2206 was found to inhibit SARS CoV-2 infection in human hepatocyte-derived cellular carcinoma cell line, Huh7 (5). Dactolisib, a PI3K inhibitor, has been demonstrated to reduce SARS CoV-2 propagation in cardiomyocytes (19). In this study, we also observed that αKG decreased the expression of other substrates of PHD2 such as HIF1α and HIF2α. A recent study has described that SARS-CoV-2 ORF3a protein induces HIF1α function, which subsequently facilitates viral infection and inflammation in patients (13). Our data therefore suggest that αKG mediated inhibition of HIFα may be involved in decreasing SARS CoV-2 replication. However, a detailed study may explore further the role αKG-PHD2-HIFα axis on SARS CoV-2 infection. In this study, we also described a significant inhibitory effect of αKG on ACE2 expression. ACE2 is a receptor for SARS CoV-2 (23). ACE2 binds to the SARS CoV-2 receptor-binding domain (RBD) and S proteins with high affinity and facilitates virus invasion into host cells (8). This may be one of the mechanisms by which αKG supplementation decreases ACE2 expression to affect the virus entry into cells and in turn inhibit SARS CoV-2 replication. We speculate that αKG mediated inhibition of pAkt may be involved in downmodulation of ACE2. Another study has described that capivasertib, a PI3K/Akt signal pathway inhibitor, decreased ACE2 expression and in turn inhibited SARS CoV-2 entry to Vero cells (20). Alternate Akt inhibitor triciribine inhibited ACE2 expression in human lung epithelial cells (6).

Our study specifically describes that supplementation of αKG, a common metabolite, significantly inhibits SARS CoV-2 replication via PHD2-pAkt axis. Dietary αKG has been used extensively for human trials. αKG (10 g/day for 2-4 months) improved appetite, increased body weight gain, and accelerated wound healing, thus improving the clinical outcome of chronic malnutrition in elderly patients (21, 22). αKG supplementation (2.5 g/day) rescued patients with chronic kidney disease from morbidities (23). Various studies, including our own works, described the safe usage of αKG in animal studies. We have described a rescue effect of this metabolite in mice from hypoxia-induced inflammation (16). αKG supplementation rescued mice from induced thrombosis and inflammation (15). Others also have used αKG for in vivo experimental therapies for manipulating multiple cellular processes related to organ development and viability of organisms, (24, 25) restriction of tumor growth and extension of survival,(26) and prevention of obesity (27).

In further investigation regarding the mechanism of αKG-mediated inhibition of SARS CoV-2, we demonstrate that αKG supplementation was unable to reduce the pAkt expression as well as SARS CoV-2 replication in PHD2-knockdown U937 monocytic cell line, suggesting a crucial role of PHD2 axis in it. On the other hand, in cells carrying PHD2, αKG inhibited pAkt and in turn reduced the pAkt-mediated SARS CoV-2 replication. Unlike αKG, Akt inhibitor TCN inhibited SARS CoV-2 replication in PHD2-knockdown cells.

SARS CoV-2 primarily affects respiratory system, more specifically the lower respiratory tract including lungs. It is now widely recognized that respiratory symptoms of COVID-19 are extremely heterogeneous, ranging from minimal to significant hypoxia with acute respiratory distress syndrome (ARDS). The severe and critical nature of the disease is a result of ARDS in lungs, characterized by elevated intravascular microthrombus, accumulation of macrophages and neutrophils, deposition of mucus, collagen and other extracellular matrix and accumulation of extracellular fluid in alveoli (28–31). This leads to a condition of reduced blood-gas barrier permeability and exchange of O_2_, causing hypoxemia (less O_2_ in blood). Patients, who survive ARDS, eventually live with fibrotic and dead lung tissues (32, 33). This is because as the body attempts to recover from ARDS, fibroblasts proliferate rapidly and lead to massive amount of extracellular matrix deposition. A large number of patients require O_2_ support even after recovery from this viral infection as the lungs of these patients are incapable of carrying out normal exchange of O_2_. Therefore, the blood concentration of O_2_ remains lower than normal in these patients. In many of these cases lungs are unable to heal, eventually leading to death of COVID-19 patients. Our study describes that supplementation with dietary αKG (400 mg/kg body wt. daily for 4-5 days of infection) significantly reduced lung inflammation (accumulation of inflammatory immune cells and pro-inflammatory cytokines, including IL6 and TNFα) and thrombus formation (platelet clots and platelet-leukocyte aggregates) in SARS CoV-2 infected hamsters and mice. Our recent works describing anti-inflammatory as well as anti-thrombotic role of αKG (15, 16) support our above observations. αKG significantly reduced the expression of pAkt and HIFα, and rescued from SARS CoV-2 infection. Interestingly, αKG supplementation also rescued the injured lungs of the infected animals from apoptotic cell death. A recent work has also described that elevated HIF2α induces ROS generation and oxidative cell death in colon cells (34). It may be possible that αKG supplementation inhibited ROS induced cell death by inhibiting HIF2α. However, this mechanism yet to be investigated in detail in our above models.

Most interesting part of our study describes that αKG supplementation rescued both hamsters and mice, although more prominent in former, from the clinical condition of hypoxemia. αKG supplementation maintained normal saturation of oxygen in circulation of the SARS CoV-2 infected animals.

Most efficient protection mechanism in viral infection is the antibody mediated defense. The CD4 T cell and B cell axis generates antibodies to neutralize the viral load and CD8 T cells eliminate the infected cells. Extensive studies describe that supplementation with limited dosage of αKG is safe in animals as well as in humans. We investigated particularly the effect of αKG on adaptive immune response in the SARS CoV-2 infected animals. Our study describes that αKG supplementation neither affect the levels of IgG against SARS CoV-2 RBD protein, nor interfered with the percentage of interferon-γ positive (IFNγ+) CD4+ and IFNγ+CD8+ T cells in infected animals. It did not alter the function of neutralization antibody against SARS CoV-2 in hamsters supplemented with αKG.

Thus, our study strongly suggests that αKG may be used as first-line treatment for COVID-19. We speculate that this metabolite may be used alone or in combination with current regimens for the disease. The employment of αKG opens up new avenues of treatment for lung inflammation and thrombosis in other such respiratory diseases as well.

## MATERIALS AND METHODS

### Ethics

Ethics approval was obtained from the Institutional Animal Ethics Committee (IAEC) (ref. no. RCB/ IAEC/2021/093), and Institutional Biosafety Committee (IBSC; ref. no. RCB/IBSC/21-22/308) of the Regional Centre for Biotechnology (RCB), Faridabad, India. Experiments using BALB/c mouse strain (RRID: IMSR_JAX_000651) and Syrian golden hamster (available form ICMR-National Institute of Nutrition, Hyderabad, India) were conducted within the guidelines of IAEC in the Biosafety level 3 (BSL3) facility of the institute.

### SARS CoV-2 infection in Vero and U937 cells

Monkey kidney epithelial cell line Vero E6 (0.1×10^6^ cells/well) was infected with P-3 Wuhan SARS CoV-2 (world reference # USA-WA-1/2020, as mentioned in our earlier work (15) at 0.01 MOI for 1 hr in Dulbecco’s Modified Eagle Medium (DMEM) with 2% FBS, supplemented with 1% non-essential amino acids. One-hour post infection, cells were supplemented with different concentrations of octyl αKG (0, 0.75, 1 and 1.5 mM, Sigma-Aldrich, USA) or Triciribine (TCN; from Selleckchem, USA) for 24 hr in BSL3 facility. The Vero E6 cell line was validated for free of mycoplasma contamination. Cell pellet was collected in Trizol. Total RNA (1 mg) was reverse-transcribed using iScript select cDNA synthesis kit (Bio-Rad, USA) as per manufacturer’s protocol, using primer 5’CGTGTCATGGTGGCGAATAAGATTAAAGGTTTATACCTTCCCAGGTAACA 3’ for the synthesis of negative strand cDNA. The cDNA was diluted in nuclease-free water (Promega, USA) and used for real-time PCR with either SARS CoV-2 or 18S rRNA specific primers, using SYBR-green mix (Bio-Rad) in an Applied Biosystems Quant Studio TM 6Flex Real-Time PCR System. The oligonucleotides used were SARS-F (5’ GTGTCATGGTGGCGAATAAG 3’) and SARS-R (5’ TCGTTGAAACCAGGGACAAGG 3’) for SARS-CoV-2, and 18S r-RNA-F (5’ GGCCCTGTAATTGGAATGAGTC 3’) and 18S rRNA (5’CCAAGATCCAACTACGAGCTT3’). The Ct value corresponding the viral RNA was normalised to that of 18S rRNA transcript. The relative level of SARS CoV-2 RNA in mock-infected samples was arbitrarily taken as 1 and that of infected samples expressed as fold-enrichment (FE). The FE was calculated as 10 logarithmic value and plotted.

SARS CoV-2 infection in U937 cells: Wild type U937 monocytic cells line or PHD2-knockdown U937 cells were transfected with a human ACE2 adenovirus (AD5CMVACE2, Viral Vector Core Web, University of Iowa, USA) with a concentration of 10 MOI for 1 hr in the presence of 8 μm polybrene. After wash with PBS, cells were kept in fresh media having 2% serum for 48 hr, then we proceeded with the SARS CoV-2 infection at 0.01 MOI for 1 hr and wash the cells and kept in fresh media for 24 hr with and without octyl αKG or TCN treatment.

### SARS CoV-2 infection in hamster and mice

We used Syrian golden hamster model of SARS CoV-2 infection as described in our previous work (15). Male hamsters of 8 weeks old were infected with one-time SARS CoV-2 via nasal route inoculation using 1×10^5^ plaque-forming units (PFU). Animals were supplemented with a human dietary grade αKG (Double Wood LLC, Philadelphia, USA) with a dose of 40 mg/100 g body wt., through oral gavage from day 1 to day 4 daily post infection. At day 5 and day 9 post infection, animals were sacrificed and blood, lung and spleen samples were harvested. Details are described in schematic Fig 3A.

The balb/c, 7-10 weeks old male mice were treated (one-time through intranasal route) with human ACE2 adenovirus (described above) using 25×10^8^ PFU for transient expression of hACE2 in respiratory track. Mice were infected with SARS CoV-2 via nasal route inoculation using 1×10^5^ PFU as described (15). Animals were supplemented with αKG with a dose of 8 mg/20 g body wt., through oral gavage from day 1 to 4 post infection. Animals were sacrificed at day 5 and day 9 post infection. Blood and lungs were collected from the animals. Details are described in schematic Fig 6A.

### PCR estimation of viral genome from lung tissue

Lung tissue sample from hamsters and mice was homogenized in Trizol using a hand-held tissue homogenizer and the total RNA extracted as per manufacturer’s protocol. RNA was reverse-transcribed using iScript™ cDNA synthesis kit (Bio-Rad) as per manufacturer’s protocol. The cDNA was used for real-time PCR for SARS CoV-2 or G3PDH, as described above. The primer used were SARS SN1-F (5’-GACCCCAAAATCAGCGAAAT -3’) and SARS SN1-R (5’TCTGGTTACTGCCAGTTGAATCTG-3’) for SARS CoV-2, and G3PDH-F (5’-GACATCAAGAAGGTGGTGAAGCA-3’) and G3PDH-R (5’-CATCAAAGGTGGAAGAGTGGGA-3’). The Ct value corresponding the viral RNA was normalised to that of G3PDH transcript. The relative level of SARS CoV-2 RNA in mock samples was arbitrarily taken as 1 and that of infected samples expressed as fold-enrichment, as described in our previous work (15).

### Confocal microscopy

Vero E6, U937 cells or U937/PHD2-KD cells from above experiment were fixed in 4% paraformaldehyde for 20 min at room temperature. After wash with 1xPBS, cells were permeabilized using 0.1% Triton X-100 followed by 2 hr blocking with 5% BSA and 5% FBS at RT. The cells were labelled with primary antibodies keep it for overnight followed by Alexa 594 conjugated secondary antibody. Further, they were stained with DAPI for 10 min and then cells were mounted with ProLong Gold anti-fade reagent. Images were captured using a Leica Confocal DMI 6000 TCS-SP8 microscope (Leica Microsystems, Wetzlar, Germany) at 63× oil immersion objective (NA 1.4) Plan Apo objectives and quantified as mentioned. Imaging was performed using Z-stacks at 0.25 μm per slice by sequential scanning and Image J Fiji software was used to obtain maximum intensity projection images. The total cell fluorescence intensity was calculated from at least 25 cells in each set per experiment and data represented as mean and standard error of mean from at least 3 independent experiments.

### Western blotting

The lyste of cells pellet or lung tissue was prepared in RIPA lysis buffer and protease-phosphatase inhibitor (Thermo Scientific Life Tech, USA) at BSL3 facility. The lysate samples were further processed for SDS-PAGE followed by immunoblotting using primary antibodies against pAkt-Ser473, Akt, pAkt1-Thr308, Akt1, HIF1α, HIF2α, β-Actin (Cell Signalling, USA), as described in our work (15).

### Immunohistochemistry, microscopy

Lungs from hamsters and mice were fixed in 10% formalin solution and processed for paraffin embedding. 3 μm thick sections of the embedded tissues were cut for staining. All samples were stained with hematoxylin and eosin; and Masson’s trichrome stains. Tissue sections from hamsters were stained for pAkt, HIF2α and Spike proteins in the lungs. Each stained section was analysed and captured at 20X magnification. Cellularity, Thrombosis score and intensity of Immunohistochemistry were quantified using ImageJ software. Cellularity was calculated as percentage of area covered by the nucleus as described in our work (15) Thrombosis and immunohistochemistry were quantified using colour deconvolution method in the software.

### Immunophenotyping

Single cell suspension was prepared from lung tissue using Liberase (100 μg/ml; Sigma-Aldrich). Chopped lung pieces were kept in Liberase for 1 hr at 37° C. After the lung chunks started to dissolve in the form of cell suspension, cells were strained using cell strainer and washed. After RBC lysis PBMCs were stained for measuring leukocytes and leukocytes-platelet aggregates. Cells were stained with anti-mouse CD45.2 APC and CD41a PE (BD Biosciences, San Jose, USA) for 30 min in dark at RT. Cells were then washed and acquired on BD FACS Verse and were analyzed with FlowJo software (Tree Star, USA).

### Cytokine assay

Cytokines such as TNFα, IL6 and IL10 were measured from mice plasma of different treatments at different day points using CBA and analysed by FCAP array software (BD Biosciences, USA).

### Platelet microparticle

Platelet poor plasma (PPP) was obtained by centrifuging Platelet rich plasma at 1000 g for 10 min. Platelet free plasma was obtained from PPP in 2 sequential steps: PPP at 2500 g for 15 min, and again at 5000 rpm for 15 min. The protocol is modified according to mouse plasma. Platelet-derived microparticles were measured using flow cytometry after labelling with antimouse CD41 PE (BD Biosciences, USA) as described in our work (15).

### Tunel assay

Assay was performed according to manufacturer’s protocol. Paraffin-embedded tissue sections were dewaxed and rehydrated using xylene and ethanol gradient. Tissues sections were then microwave irradiated in 0.1 M citrate buffer, pH 6.0 for 10 min. 50 μl TUNEL reaction mixture was homogeneously spread over the tissue section and incubated in a humidified atmosphere for 60 min at 37° C in the dark. Sections were washed and mounted with mounting media prolong gold. Image was captured in fluorescence microscopy with an excitation wavelength in the range of 520-560 nm and detection in the range of 570-620 nm.

### SARS CoV-2 RBD antibody measurement

The binding-antibody response to SARS CoV-2 post-infection was measured using ELISA based platform as described earlier (*35, 36*). The antibody response was measured against RBD protein post virus challenge at different time points. Soluble RBD protein was coated on Maxisorp plates (Nunc) with 2μg/ml of purified protein and incubated with three-fold serially diluted sera (with 1:5 as starting dilution) in dilution buffer (1:5 times dilution of blocking buffer). The plates were incubated at room temperature (RT) for 1 h and then washed three times with washing buffer (PBS + 0.1 % tween 20) and incubated with HRP conjugated anti-hamster IgG antibody at 1:10,000 dilution for another 1 h and washed four times with the washing buffer and incubated further with 100 μl of TMB substrate (Thermo Fisher Scientific), The reaction stopped with 1N H_2_SO_4_ solution. Sera end point titers were calculated as the reciprocal of serum dilution giving OD 450 nm readings higher than lowest dilution of the placebo or control arm + two times standard deviations.

### T cell activation assay

Splenocytes were isolated from the hamster spleen. Spleen was crushed using frosted slide and single cell suspension was prepared using wire mesh. After RBC lysis from single cell suspension, splenocytes (1 million cells from each animal) was stimulated with PMA (phorbol 12-myristate13-acetate; 50 ng/ml; Sigma-Aldrich), ionomycin (1.0 μg/ml; Sigma-Aldrich) for 6 hr. Cell surface staining with anti-mouse CD4 PE and CD8 APC (BD Biosciences, USA) was carried out for 30 min in dark at RT. Cells were then fixed in IC fixation buffer (Thermo Fisher Scientific, USA) for 30 min each. Intracellular anti-mouse IFN-γ APC Cy7 (BD Biosciences, USA) staining was then carried out for 30 min in dark at RT. Cells were acquired and analysed as described above.

### Neutralizing antibody measurement

Vero E6 cells were maintained in DMEM medium supplemented with 10% FBS, 100 U/ml of penicillin-streptomycin and 1% non-essential amino acids. The assay was performed in duplicate using 24-well tissue culture plates, in a biosafety level 3 facility. Serial dilutions of each plasma sample were incubated with 30-40 PFU of virus for 1 hr at 37°C. The virus-serum mixtures were added to Vero E6 cell monolayers and incubated for 1 hr at 37°C in 5% CO2 incubator. The cell monolayer was then overlaid with 2% CMC in cell culture medium and incubated for 2 days, at which time the plates were fixed in 3.7% formaldehyde and stained with crystal violet dye. PRNT50 represents antibody concentration at which 50% reduction in plaque formation occurs. PRNT50 was calculated with the help of non-linear curve using GraphPad Prism software.

### SpO_2_ measurement

Oxygen saturation (SpO_2_) in the blood was measured in both mice and hamsters from the above experiments. The Mouse STAT Jr. pulse oximeter by Kent Scientific Corporation, Torrington, USA was used for the measurement of SpO_2_ in awake animals. Paw of the animals was captured in the sensor.

### Statistical analysis

Data from at least three experiments are presented as mean ± SEM (standard error of the mean). Statistical differences among experimental sets with normally distributed data were analyzed by using one-way or two-way ANOVA followed by Bonferroni’s correction for multiple comparison. Kruskal Wallis test followed by Dunn’s multiple comparison post-test was used for non-normally distributed data. D’Agostino-Pearson Test was used to check for normal distribution of data. Graph Pad Prism version 8.0 software was used for data analysis and P-values.

## Supplementary Materials

**Fig.S1.** Viral replication kinetics and cell death assay at 12, 24 and 36 hr after infection *in vitro*.

**Fig S2.** Densitometry analysis of all western blot data.

**Fig. S3.** Body weight of the hamsters during infection.

**Fig. S4.** Blood counts of hamsters from treated groups.

**Fig.S5.** More representative images of TUNEL assay in the lungs of infected hamsters to assess apoptotic cells.

**Fig.S6.** Leukocytes percentage and Platelet-leukocytes aggregates in blood of hamsters.

**Fig.S7.** Gating strategy for interferon-γ positive (IFNγ+) CD4+ and IFNγ+CD8+ T cells.

## Acknowledgements

This study is supported by grants: BT/PR22881 and BT/PR22985 from the Department of Biotechnology (DBT), Govt. of India; and CRG/000092 from the Science and Engineering Research Board, Govt. of India to PG. Authors thank to Dr. Arundhati Tiwari of Regional Centre for Biotechnology, Faridabad, India, for English editing.

## Authors contributions

SA and SK designed and performed all experiments and analyzed data. TRA and GJ designed and performed in vitro and hamster infection experiments and measured viral load and performed neutralization assay. NS has performed western blot analysis. AS has performed confocal microscopy study. OS performed mice infection experiment. TS has conceptualized and performed antibody measurement assay and analyzed data and edited manuscript. PS has performed antibody measurement assay. SV and MS have conceptualized infection studies and edited manuscript. PG designed and supervised the study, conceptualized the approach, designed the experiments, analyzed the data, and wrote the manuscript. All authors read, edited and approved the final manuscript.

## Data and materials availability

All data are available in the main text or the supplementary materials.

## Online Supplementary Information

### Supplementary Figures

**Fig.S1.**
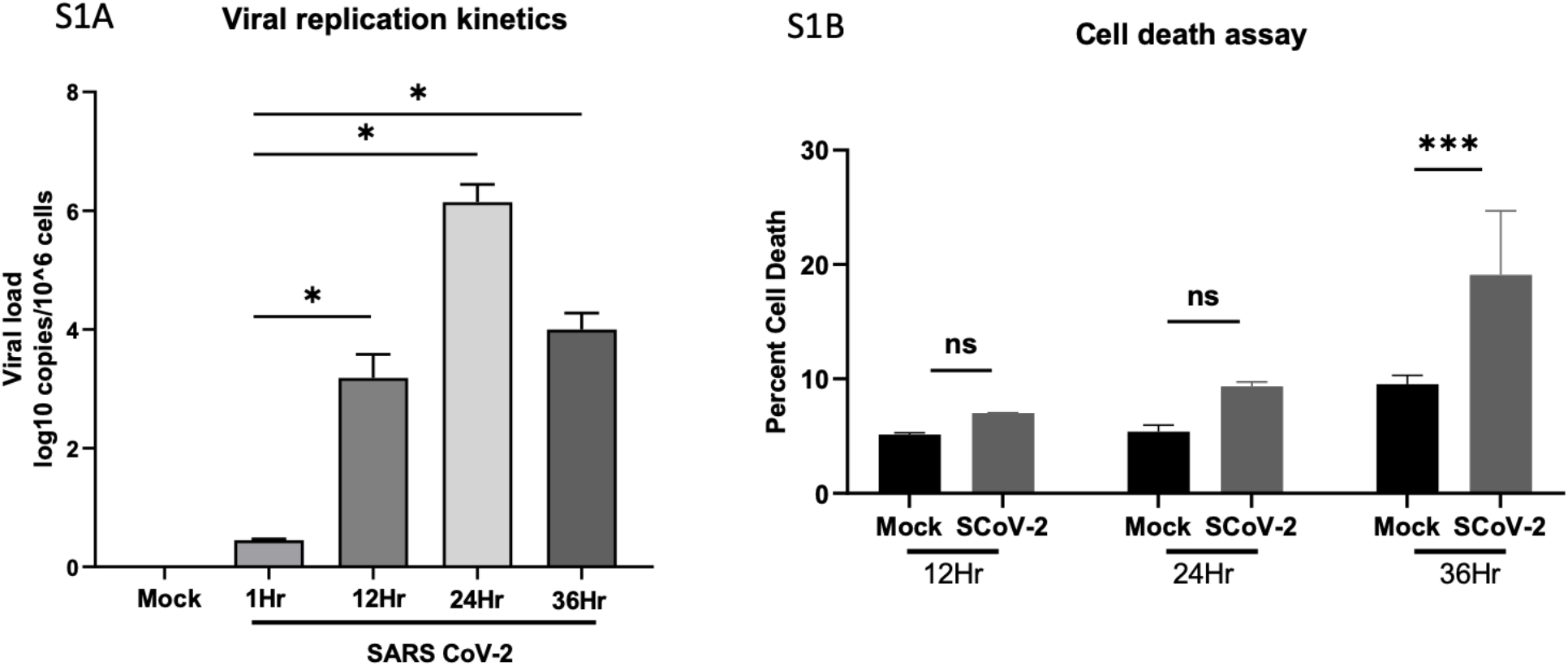
Viral replication kinetics and cell death assay: (A) Viral replication (measured by RT-PCR) increased at 12hr and 24 hr post infection. And further reduced at 36 hr post infection. Data are mean ± SEM from 3 independent experiments. (B) Cell death increased time dependently in SARS CoV-2 infected cells. Measured using Propidium iodide staining. Data are mean ± SEM from 3 independent experiments. Mann-whitney unpaired t test was used to compare between the groups (\**P*<0.05, \*\*\**P*<0.001 and \*\*\*\**P*<0.0001).

**Fig S2.**
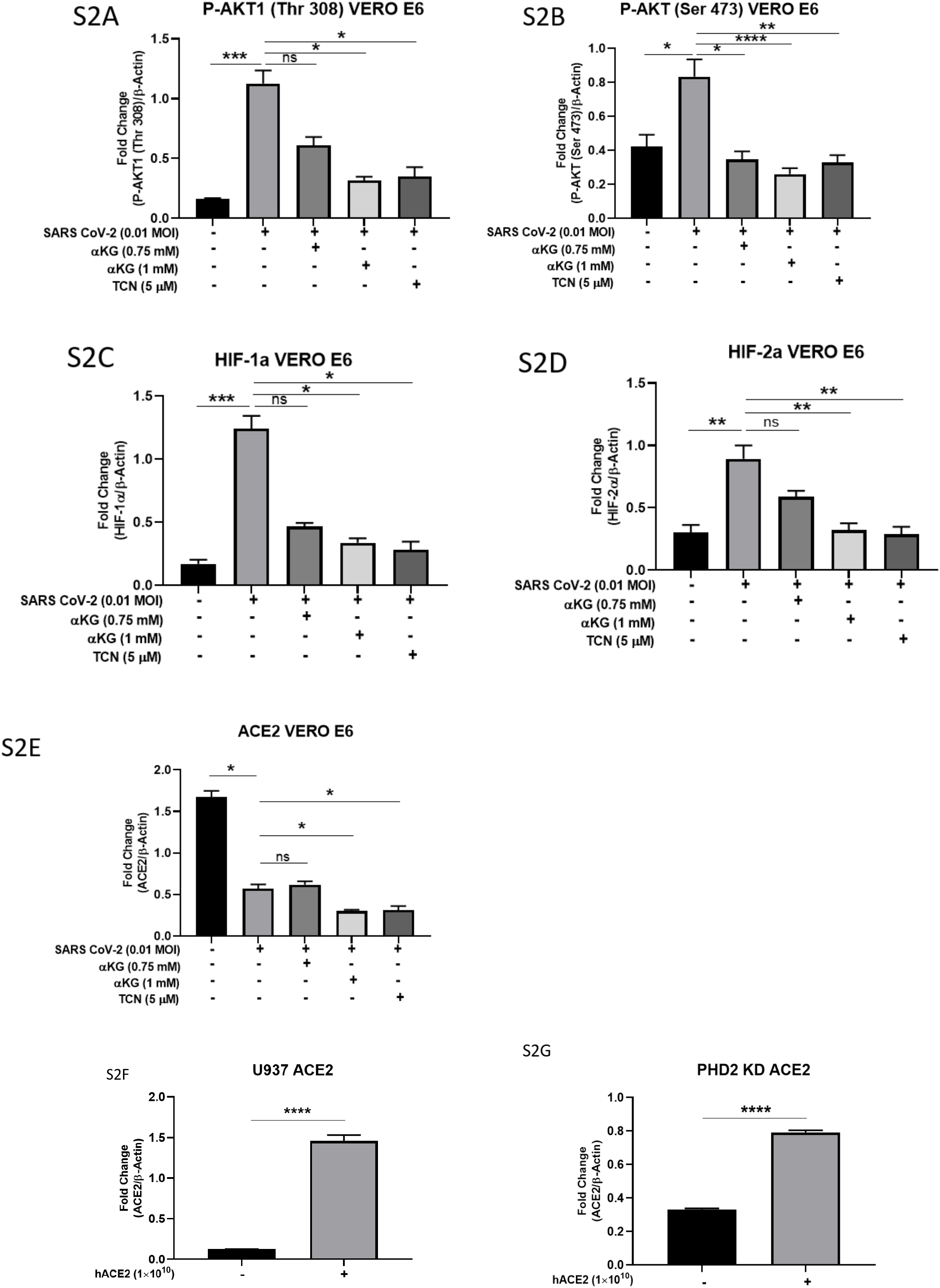
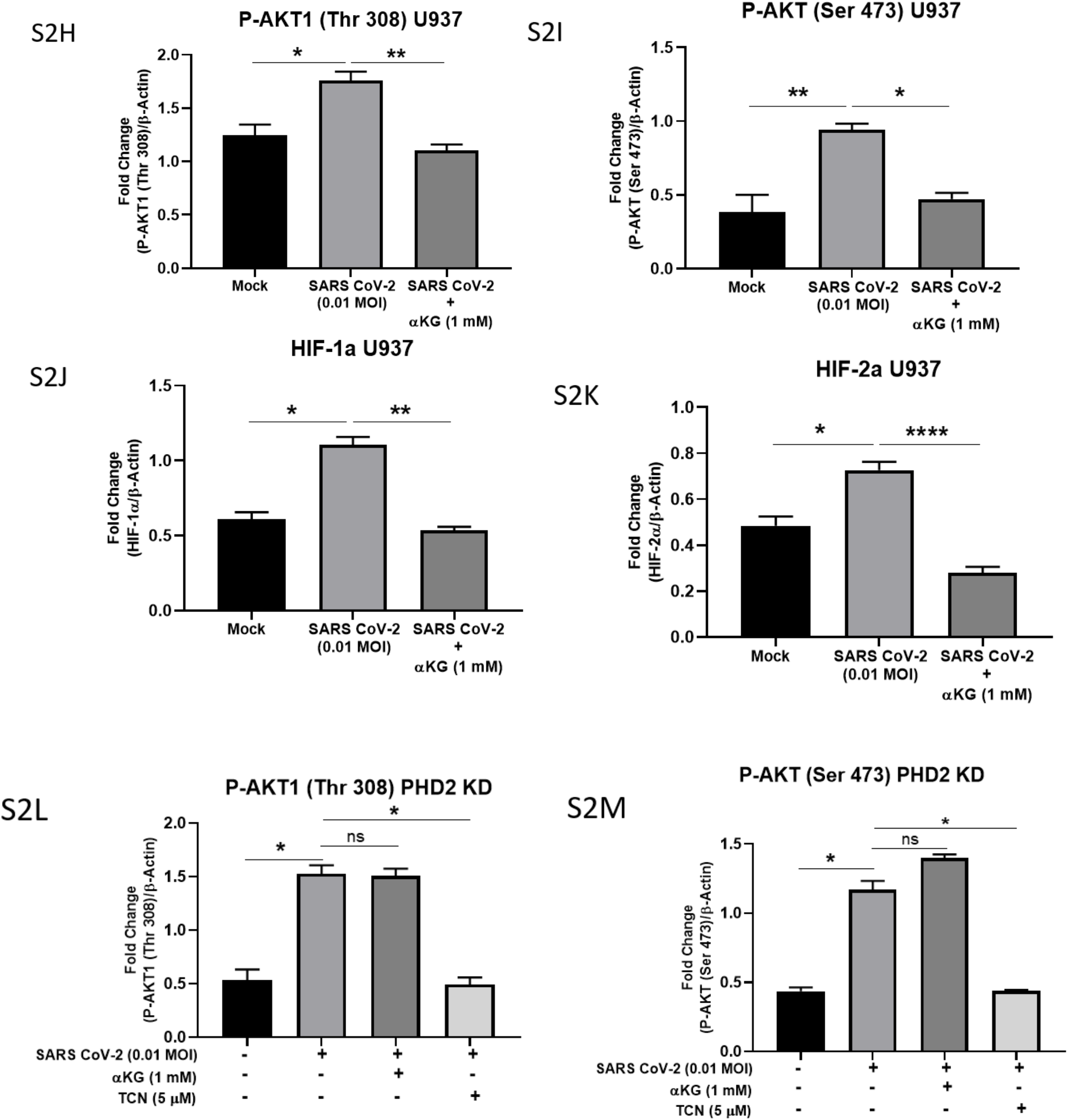
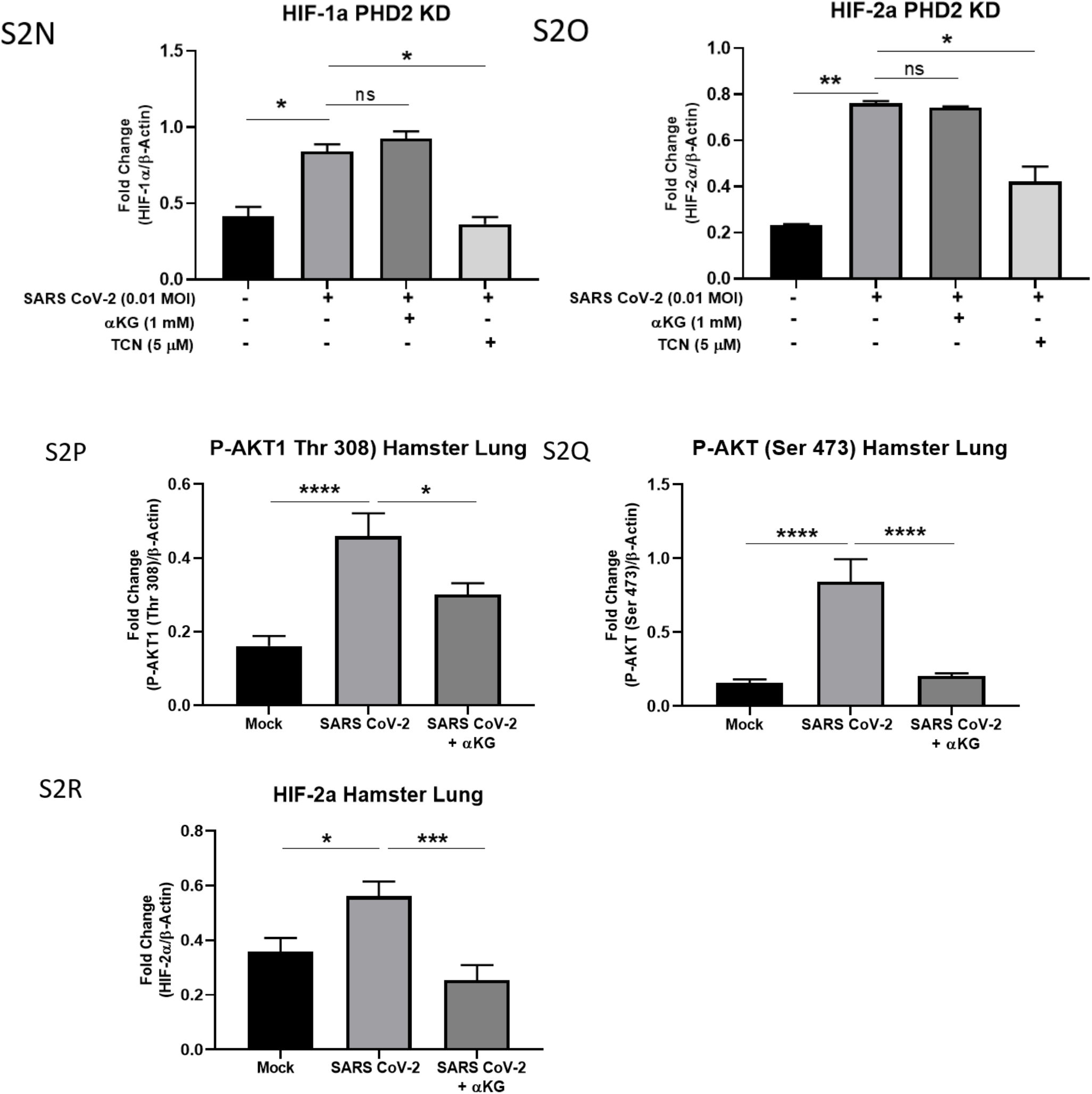
Densitometry of western blots. Densitometric evaluation of the protein bands was performed using the Alpha imager software for western blots. Densitometry related to western depicted in main Fig. 1D (Fig. S2A-E), Fig. 2A-B (Fig. S2F-G), Fig. 2I (Fig. S2H-K), Fig. 2J (Fig. S2L-O), Fig. 3H (Fig. S2P-R). Bar represents mean ± SEM from triplicate blots. Kruskal Wallis test used to compare between the groups, \**P*<0.05, \*\**P*<0.01, \*\*\**P*<0.001, **** *P*<0.0001, ns=non-significant.

**Fig. S3.**
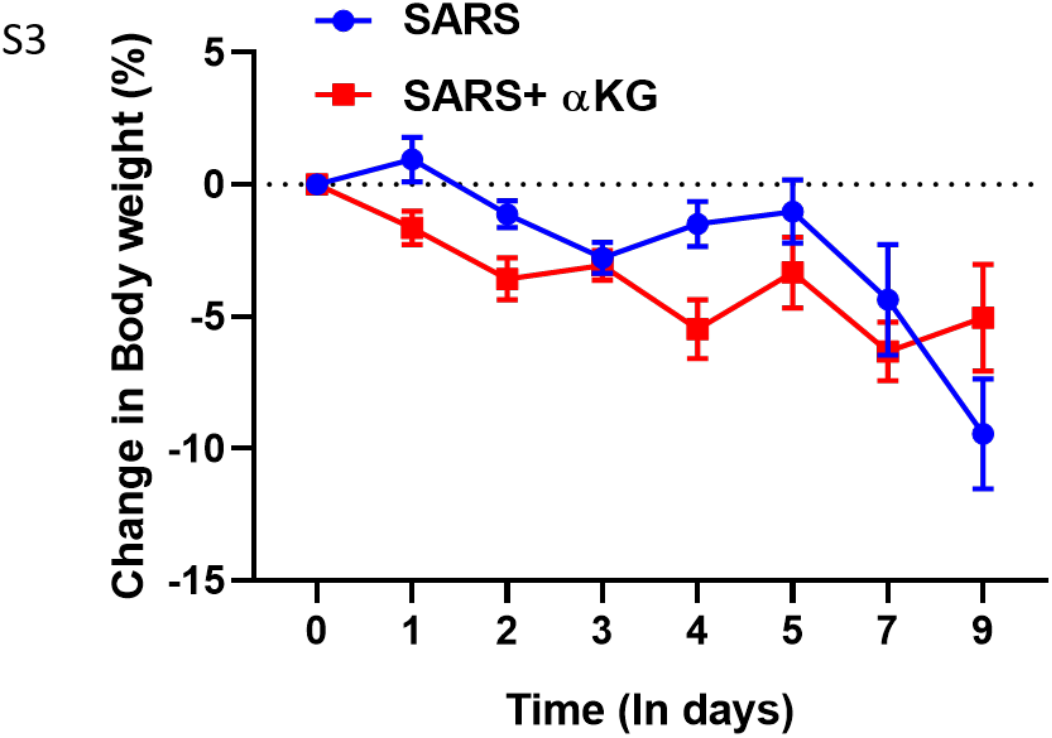
Body weight of the Hamsters. Body weight of animals were measured till day of sacrifice, starting from the day of infection. Data is from 10-15 animals in each group represented as mean ± SEM.

**Fig. S4.**
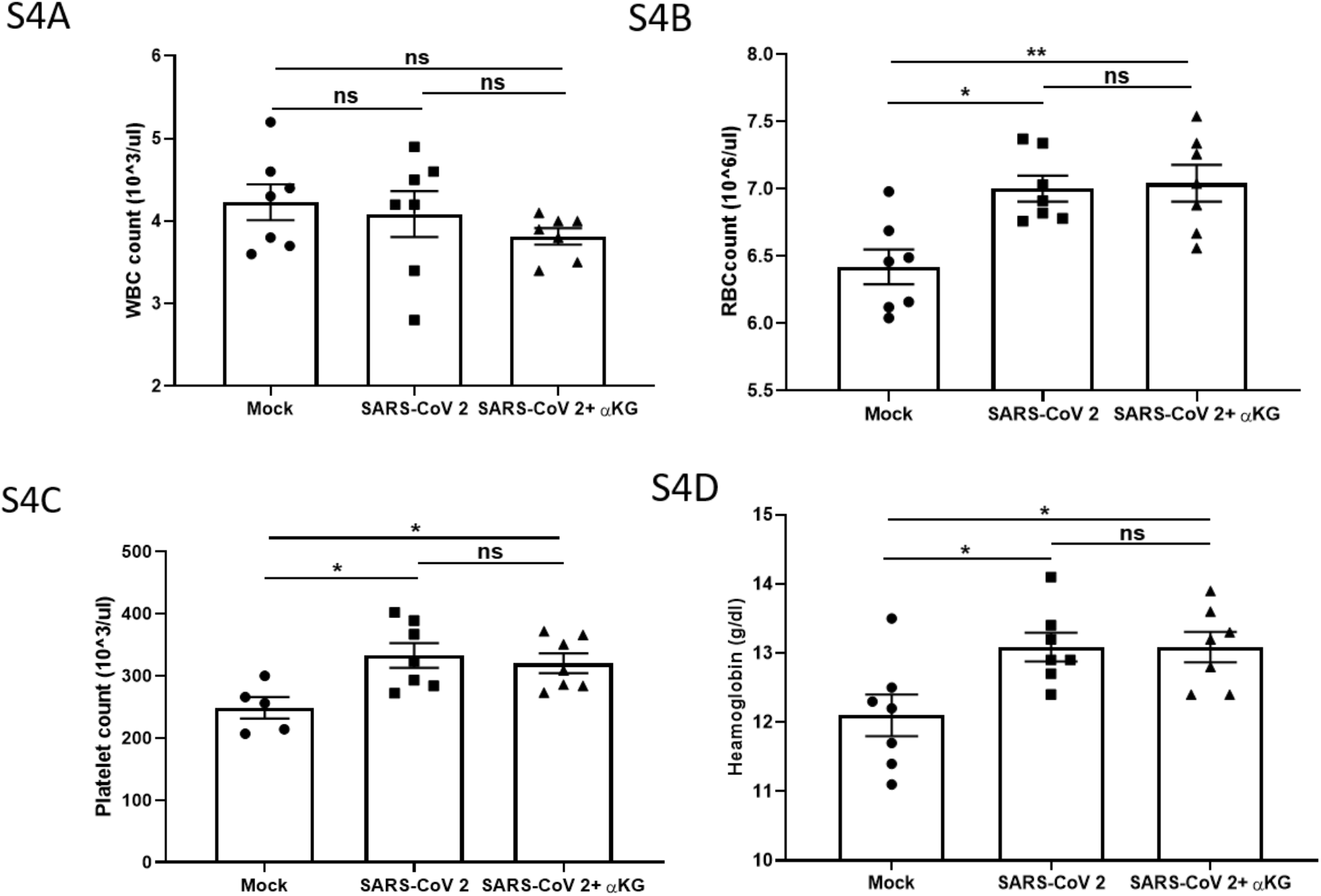
Hamster blood counts. Count of (A) WBC, (B) platelets, (C) RBC and (D) Haemoglobin determined from SARS CoV-2 Wuhan P8 strain infected hamsters blood with or without αKG treatment at 5dpi (as mentioned in Fig. 6), using Nihon Kohden’s CelltacF haematology analyser. Data are mean ± SEM. One way ANOVA test was used to compare between the groups, (\**P*<0.05, \*\**P*<0.01 ns=non-significant).

**Fig.S5.**
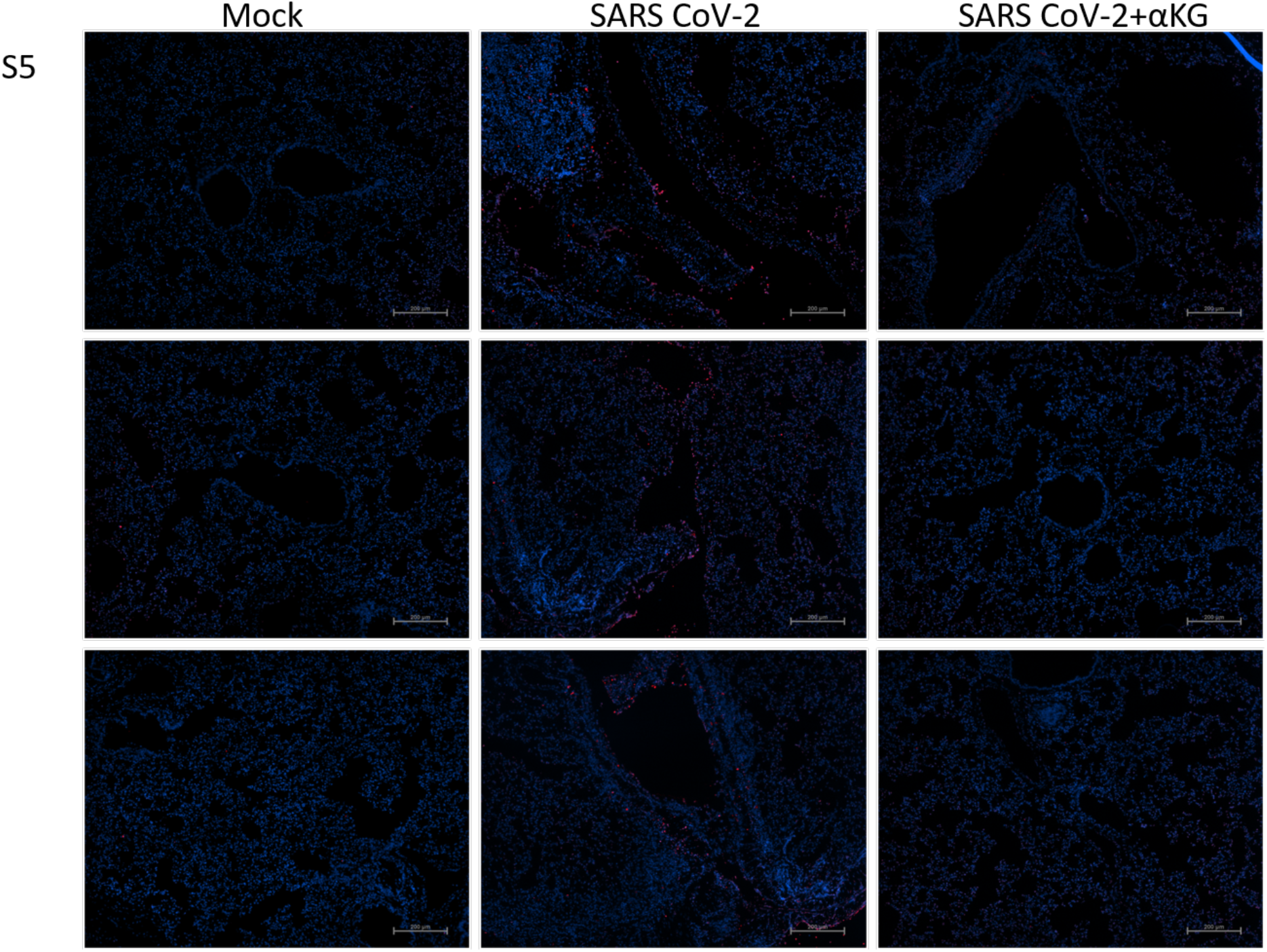
More representative images of TUNEL assay in the lungs of infected hamsters to assess apoptotic cells as mentioned in Fig. 4I-J. Less number of cells in αKG treated hamster lungs at 5 dpi.

**Fig.S6.**
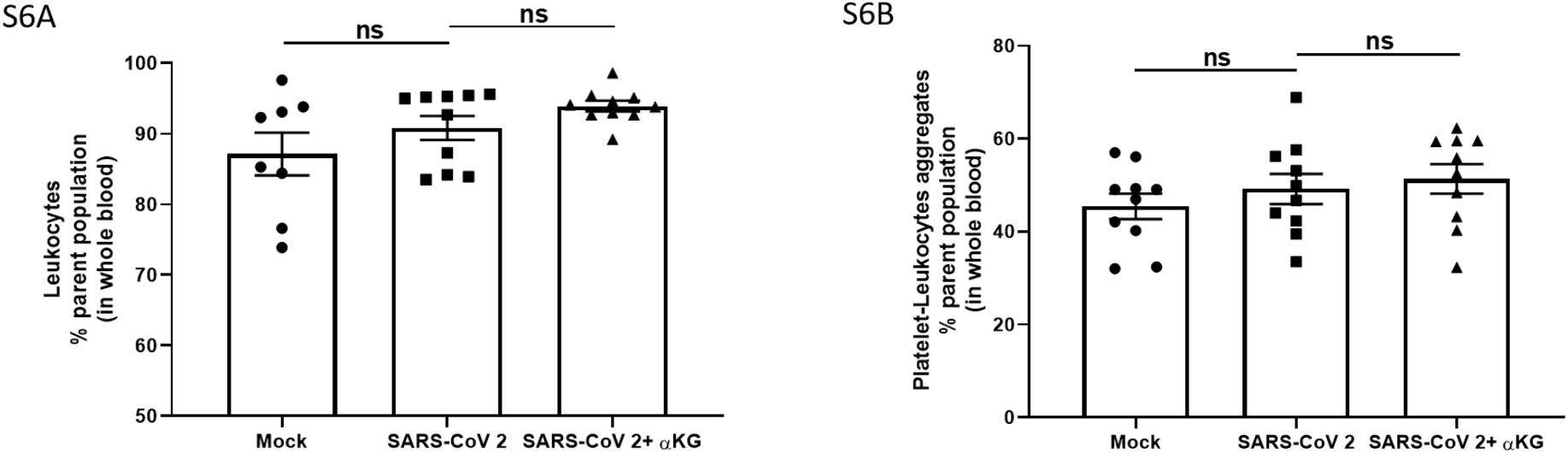
Leukocytes percentage and Platelet-leukocytes aggregates in blood. (A) number of Leukocytes and (B) Number of platelet-leukocytes aggregates measured from whole blood using flow cytometry in SARS CoV-2 infected hamsters with and without α-KG treatment at 9 dpi. Data is mean + SEM. One way ANOVA, Bonferonni’s post-test, ns= non significant.

**Fig.S7.**
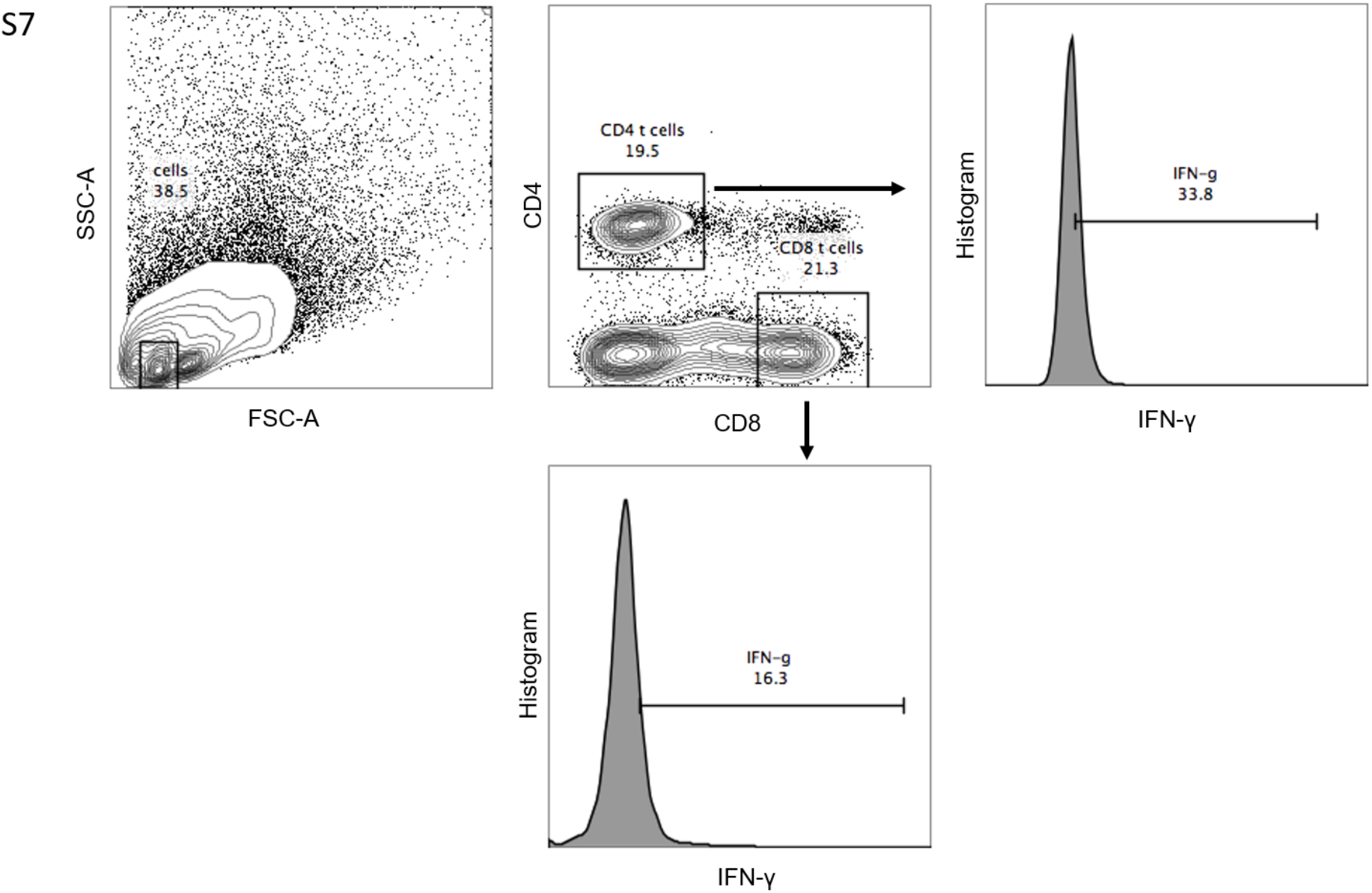
Gating strategy for interferon-γ positive (IFNγ+) CD4+ and IFNγ+CD8+ T cells.

